# Emergence of metabolic coupling to the heterotroph *Alteromonas* promotes dark survival in *Prochlorococcus*

**DOI:** 10.1101/2024.05.01.591941

**Authors:** Allison Coe, Rogier Braakman, Steven J. Biller, Aldo Arellano, Christina Bliem, Nhi N. Vo, Konnor von Emster, Elaina Thomas, Michelle DeMers, Claudia Steglich, Jef Huisman, Sallie W. Chisholm

## Abstract

*Prochlorococcus* is found throughout the euphotic zone in the oligotrophic open ocean. Deep mixing and sinking while attached to particles can, however, transport *Prochlorococcus* cells below this sunlit zone, depriving them of light for extended periods of time. Previous work has shown that *Prochlorococcus* by itself cannot survive extended periods of darkness. However, when co-cultured with a heterotrophic microbe and subjected to repeated periods of extended darkness, *Prochlorococcus* cells develop an epigenetically inherited dark-tolerant phenotype that can survive longer periods of darkness. Here we examine the metabolic and physiological changes underlying this adaptation using co-cultures of dark-tolerant and parental strains of *Prochlorococcus*, each grown with the heterotroph *Alteromonas* under diel light:dark conditions. The relative abundance of *Alteromonas* was higher in dark-tolerant than parental co-cultures, while dark-tolerant *Prochlorococcus* cells were larger, contained less chlorophyll, and were less synchronized to the light:dark cycle. Meta-transcriptome analysis revealed that dark-tolerant co-cultures undergo a joint change, in which *Prochlorococcus* undergoes a relative shift from photosynthesis to respiration, while *Alteromonas* shifts towards using more organic acids instead of sugars. Furthermore, the transcriptome data suggested enhanced biosynthesis of amino acids and purines in dark-tolerant *Prochlorococcus* and enhanced degradation of these compounds in *Alteromonas*. Collectively, our results demonstrate that dark adaptation involves a strengthening of the metabolic coupling between *Prochlorococcus* and *Alteromonas*, presumably mediated by an enhanced, and compositionally modified, carbon exchange between the two species.

## Introduction

*Prochlorococcus* is the most abundant photoautotrophic organism of the tropical and subtropical ocean, with an estimated global population of *∼*3x10^27^cells (1). These small non-motile cells are found throughout the euphotic zone (∼0-200 m), reaching densities of ∼10^5^ cells mL^-1^ in surface waters and up to 10^4^ cells mL^-1^ at the base of the euphotic zone (2). *Prochlorococcus* also occurs below the euphotic zone - presumably due to deep vertical mixing or aggregation with sinking particles - and have been detected in metagenomes as deep as the bathypelagic zone (∼1000-4000m) (3–5), and have also been found at densities as high as 10^2^-10^4^ cells mL^-1^ in the mesopelagic zone (∼200-1000m) (6,7). However, it is unclear whether *Prochlorococcus* cells in the aphotic zone are alive or dead. If viable, they would represent a sizable unaccounted fraction of the global *Prochlorococcus* population that could not only affect global biogeochemical processes, but also represent a large genetic reservoir shaping *Prochlorococcus* ecology and evolution.

We have been studying whether and how *Prochlorococcus* survives extended periods of darkness to better understand this phenomenon. We previously found that axenic cultures cannot survive more than a day in the dark, but when co-cultured with heterotrophic bacteria of the *Alteromonas* genus, *Prochlorococcus* cells can survive up to 11 days of darkness (7). Moreover, axenic cells can survive 3 days of darkness when the media is amended with both a reactive oxygen species scavenger (pyruvate) and an organic carbon substrate (glucose) (7), suggesting that heterotrophic metabolism of organic carbon obtained from its co-culture partner could be important for dark survival of *Prochlorococcus*. Indeed, *Prochlorococcus* is capable of assimilating diverse organic compounds (8–12) and it has been estimated that cells at the bottom of the euphotic zone obtain >80% of carbon from organic compounds (13), supporting the potential importance of heterotrophic metabolism in *Prochlorococcus*. We have shown further that when *Prochlorococcus* cells are subjected to repeated episodes of extended darkness, only co-cultures with heterotrophic bacteria exhibit significantly faster growth recovery relative to parental cells, producing a non-genetic, heritable dark-tolerant phenotype (14). The emergence of dark-tolerance in *Prochlorococcus* is a facilitated adaptation as it depends on the presence of the heterotroph and does not occur in axenic cultures (14).

Here, we investigate the metabolic basis of dark-tolerance in *Prochlorococcus*, and begin to unpack the nature of its metabolic interactions with *Alteromonas*. For this purpose, we compare the population dynamics and cell cycle in co-cultures of dark-tolerant *Prochlorococcus* and *Alteromonas* with those in parental co-cultures that have not been adapted to extended darkness. Furthermore, we analyze the meta-transcriptome of *Prochlorococcus* and *Alteromonas* in dark-tolerant and parental co-cultures to assess changes in important metabolic processes such as the electron transport chain, Calvin Cycle, EMP glycolysis, and the TCA cycle. The results suggest that *Prochlorococcus’* adaptation to survive periods of extended darkness is due to a decreased use of photosynthesis and an increased use of exogenous organic carbon, mediated by cross-feeding with the heterotroph *Alteromonas*.

## Materials and Methods

### Cultures

Co-cultures of *Prochlorococcus* NATL2A and *Alteromonas macleodii* MIT1002 were prepared as previously reported (14) and were grown in a 13:11 light dark cycle with simulated dawn and dusk (15) at 37 *μ*mol photons m^−2^ s^−1^ in 24°C. Exponentially growing cultures were placed into a 24°C dark incubator at “sunset” for a total of 83 h (∼3.46 d) of darkness (referred to as 3 days of darkness hereafter). As described in Coe et al (2021), after 3 days of dark, cultures were shifted back into the 13:11 light dark incubator at “sunrise” and recovery was monitored via bulk chlorophyll fluorescence (10AU model, Turner Designs, Sunnyvale, California) and flow cytometry (see below). Once recovered cells reached late exponential growth, they were transferred into fresh media and the process was repeated 6 more times. After the 7th transfer, cells were moved to continuous light at equivalent integrated photons day^-1^ for 7 additional transfers (no extended darkness) to determine if the phenotype remained stable and heritable. On the 15th transfer, cells were returned to the 13:1l light:dark incubator and subjected to 3 days of extended darkness as described above. Measurements and sampling during dark periods were performed under green light (7,16).

### Flow cytometry and Cell Cycle analysis

Using sample preparation methods previously described (14), *Prochlorococcus* cell abundance samples were processed on a Guava 12HT flow cytometer (Luminex Corp., Austin, TX, U.S.A.) by exciting cells with a blue 488 nm laser and analyzing for chlorophyll fluorescence (692/40 nm), 1x SYBR Green I (Invitrogen, Grand Island, New York, U.S.A.) stained DNA fluorescence content (530/40 nm), and cell size (forward scatter). Calculations for relative cell size and chlorophyll per cell were performed by normalizing to 2 μm diameter beads (catalog no. 4500-0025, Guava easyCheck kit, Luminex Corp.) as previously described (7). Flow cytometry was analyzed using FlowJo version 10.7.1 (Flowjo LLC, BD Life Sciences, Ashland, OR, USA) and cell cycle analyses were performed using ModFit LT version 5.0 (Verity Software House, Topsham, ME USA).

### Transcriptomic sampling

Samples were taken from exponentially growing cells in standard 13:11 light:dark conditions, 2 days prior to placement into extended darkness on the seventh transfer. The cultures were sampled across one complete diel cycle at 0, 4, 8, 16, 20, and 24 h by removing 8 mL of culture and placing it immediately into 24 mL of 4°C RNALater. Samples were incubated at 4°C for up to 5 days, filtered onto 25mm 0.2µm Supor filters (Pall, Port Washington, NY), and stored at −80°C until RNA extraction as previously described (17).

### RNA extraction and RNA-Seq library construction

As described in Biller et al (2018), cell biomass filters were incubated in 10 mM Tris (pH 8), 20,000 U of Ready-Lyse lysozyme (Epicentre, Madison, WI, USA), and 40 U of SUPERase-In RNase inhibitor for 5 min at room temperature to extract total RNA. Following the manufacturer’s instructions for the mirVana microRNA (miRNA) extraction kit (Ambion, Carlsbad, CA, USA), RNA was extracted.

Ribosomal RNA was depleted using the RiboZero kit (Illumina, San Diego, CA, USA) and strand-specific transcriptome sequencing (RNA-Seq) libraries were constructed using the KAPA RNA HyperPrep kit (Illumina, San Diego, CA, USA). Sequencing was carried out on an Illumina NextSeq 500 instrument at the BPF Next-Gen Sequencing Core Facility at Harvard Medical School, with a High-Output 75-cycle kit to obtain Single-Read 75bp reads. Libraries yielded ∼1.3 to 5 million reads (average, 2.3 million) per library (Table S1).

### RNA-seq statistical analysis

Adapters and low-quality sequence regions were removed with BBDuk (V38.16)(18), with the settings ktrim=r, k=23, mink=11, hdist=1, qtrim=rl, trimq=6. Trimmed reads were aligned to the *Prochlorococcus* NATL2A (accession # NC_007335.2) and *Alteromonas macleodii* MIT1002 (accession # NZ_JXRW01000001.1) genomes as in Biller et al. 2018 (Table S1). The number of reads that aligned to each annotated ORF in the “sense” orientation was determined using HTSeq (V0.11.2) (19), with default parameters and the “nonunique all” option (Table S1).

Genes which displayed differential expression were identified using the DESeq2 R package (V1.24.0) (20). Reads that mapped to the *Prochlorococcus* and *A. macleodii* MIT1002 genomes were analyzed separately (Table S1). Tests for differential expression were then performed on each pairwise comparison of interest for *Prochlorococcus* and *A. macleodii* MIT1002 with the Wald test, using a negative binomial generalized linear model. P-values were adjusted for multiple testing with the Benjamini–Hochberg procedure, and genes with an adjusted p-value of <0.01 were considered to be significantly differentially abundant (20). Results were visualized using ggplot2 (21).

### Cell Cycle Periodicity

Genes that showed periodic expression over the diel cycle in *Prochlorococcus* (Table S3) were detected with the R package RAIN (22), using the default workflow and a corrected p-value of < 0.1.

## Results and Discussion

### Obtaining the dark-tolerant phenotype

To study the metabolic basis of dark-tolerance, we first generated the dark-tolerant phenotype as described previously (14) by subjecting exponentially-growing cultures of *Prochlorococcus* NATL2A co-cultured with *Alteromonas* to 3 days of darkness, and then returned them to a 13:11 h light:dark cycle. Once cultures resumed growth and reached late exponential growth phase, they were transferred into fresh media and the process was repeated until dark-tolerance – defined here by the ability to resume growth within 1-2 days after dark exposure – was achieved. Dark-tolerance was reproducibly achieved after three consecutive transfers and remained stable over many generations (at least 15 transfers). The dark-tolerant cells grew more slowly than the parental cells under light:dark conditions (0.31 day^-1^ vs 0.50 day^-1^), but they had the same growth rate in constant light (0.61-0.63 day^-1^; Fig. 1 A & B), suggesting that changes in the entrainment of *Prochlorococcus* to the light:dark cycle may play a role in dark-tolerance.

**Figure 1.**
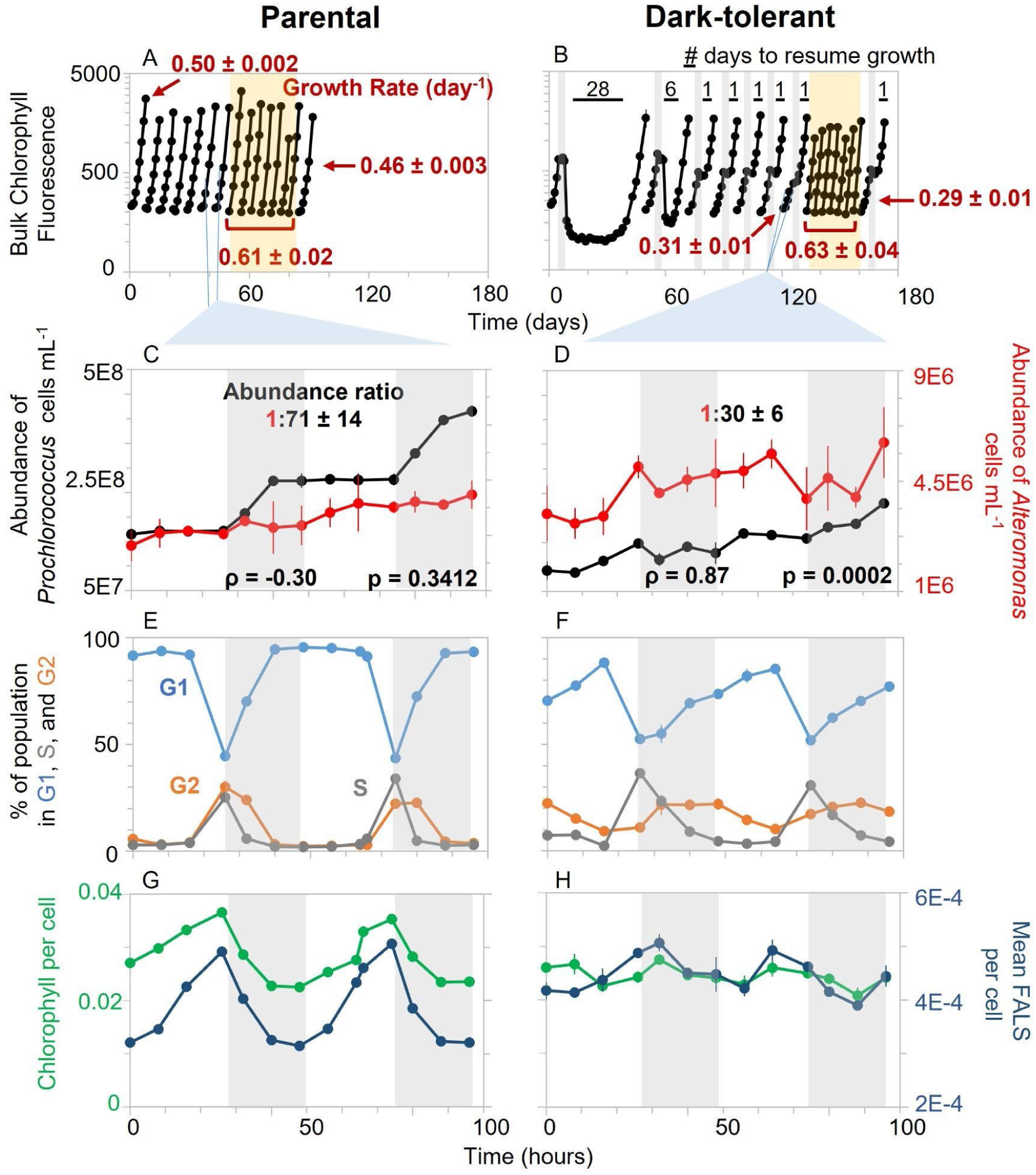
Population dynamics and cell cycle progression of parental and dark-tolerant co-cultures of *Prochlorococcus* NATL2A and *Alteromonas* grown on a 13:11 h light:dark cycle. (A) Parental co-cultures were grown under standard 13:11 h light:dark conditions. When cells reached late-exponential growth phase, they were transferred to fresh media and this process was repeated several times. After seven transfers, cells were moved into continuous light (yellow shading) for seven transfers, at an equivalent amount of photons day^-1^, before returning to light:dark conditions. (B) To establish the dark-tolerant phenotype, co-cultures of *Prochlorococcus* and *Alteromonas* were subjected to 3 days of extended darkness halfway their exponential growth phase (vertical gray bars in B) and then allowed to recover under standard 13:11 light:dark growth conditions. The time required to resume growth after dark exposure is indicated above the black horizontal bars. Again, after seven transfers, cells were moved into continuous light (yellow shading) for seven transfers, before returning to light:dark conditions. (C-H) The parental and dark-tolerant co-cultures were monitored over a 48 h period (indicated by the light blue lines below A and B) under 13:11 light:dark conditions to study (C,D) the population dynamics of *Prochlorococcus* and *Alteromonas*, (E,F) cell cycle phase analysis of the *Prochlorococcus* population (percentage in G1, S, and G2 phase), and (G,H) relative bulk chlorophyll fluorescence and mean forward angle light scatter (FALS, a proxy for cell size) per *Prochlorococcus* cell. In (C) and (D), we calculated Pearson’s correlation coefficient (ρ) between the short-term increments in population abundance of *Prochlorococcus* and *Alteromonas*, and their significance (*p*), to assess population synchrony. Gray shading in (C-H) indicates the 11 h night period on the diel cycle. Error bars for triplicate cultures are included, but may not always extend beyond the data point.

### Cell cycle and population dynamics

To understand how dark-tolerance affects growth on a light:dark cycle, we compared cell division and cell cycle progression of dark-tolerant and parental *Prochlorococcus* in co-cultures with *Alteromonas* using flow cytometry. In the parental cultures, cell division of *Prochlorococcus* was tightly synchronized with the light:dark cycle, occurring only during the first half of the dark period (Fig. 1 C, black line), as has been previously reported (15,23). In contrast, in dark-tolerant cultures, *Prochlorococcus* cells divided both in the light and in the dark, suggesting a loosening of the coupling of cell division to the light:dark cycle (Fig. 1 D, black line). Accordingly, parental cultures displayed rapid progression through the S (chromosomal replication) and G2 (cell division) phases of the cell cycle at dusk (Fig. 1E), whereas S and G2 phases extended over the entire night period in the dark-tolerant cultures, with fewer cells in G1 phase throughout (Fig. 1 F). Finally, forward angle light scatter per cell (FALS) - a proxy for cell size - and chlorophyll fluorescence per cell were both tightly entrained to the light:dark cycle in parental lines, but less so in the dark-tolerant cultures (Fig. 1 E & F). Dark-tolerant cells were on average larger than parental *Prochlorococcus*, consistent with the increased proportion of cells in the G2 phase throughout the light:dark cycle. Finally, dark-tolerant cells had a lower average chlorophyll fluorescence than parental cells, suggesting they may have lower light-harvesting capability. Together these observations further support the hypothesis that the emergence of dark-tolerance in *Prochlorococcus* involves a loosening of the coupling between the daily light:dark cycle and cell cycle progression and a decreased reliance on photosynthesis.

To understand the decreased entrainment of dark-tolerant *Prochlorococcus* to the light:dark cycle in the context of its interactions with *Alteromonas*, we examined features of both partners under the two conditions. The relative abundance of *Alteromonas*, as measured by the *Alteromonas*:*Prochlorococcus* ratio, increased from 1:71 in parental co-cultures to 1:30 in the dark-tolerant co-cultures (Fig. 1 C & D). Since *Alteromonas* growth depends fully on *Prochlorococcus* exudates, this suggests that a greater proportion of *Prochlorococcus* exudates was available to *Alteromonas* in dark-tolerant relative to parental co-cultures. Further, unlike in the parental co-cultures, changes in the population abundances of *Alteromonas* and *Prochlorococcus* over time were highly correlated in the dark-tolerant co-cultures (Pearson’s correlation: ρ = 0.87, p <0.005) (Fig. 1 C & D), indicating an enhanced synchrony between both taxa. This suggests that the decreased entrainment of *Prochlorococcus* with the light:dark cycle is linked to an increased metabolic coupling between *Prochlorococcus* and *Alteromonas* in the dark-tolerant co-cultures.

### Transcriptional patterns in Prochlorococcus

To gain additional insights into the metabolism of the dark-tolerant phenotype, we compared the transcriptomes of parental and dark-tolerant *Prochlorococcus* cells, when grown in co-culture with *Alteromonas*, every 4 h over a full light:dark cycle using RNA-Seq (Table S2). While nearly all *Prochlorococcus* genes retained periodicity in dark-tolerant cells (Table S3), the vast majority of genes showed significant differences in expression relative to parental cells, with the largest magnitudes of expression differences occurring at night (Fig. S1, Text S1). Below we highlight key differences in transcript expression patterns of dark-tolerant cells relative to parental cells.

#### Regulation of circadian rhythm

Given the decoupling of the cell cycle from the light:dark cycle (Fig. 1) and maintenance of periodic gene expression in dark-tolerant cells (Table S2), we first examined the expression patterns of genes related to the circadian clock. *Prochlorococcus* has a partial KaiBC circadian oscillator which must be reset each morning to sustain periodicity, likely through interactions that sense the cellular redox state (24–26). It uses a two-component regulatory system, SasA-RpaA, to transmit the clock’s output to the rest of the genome, ultimately driving genome-wide transcription rhythms (27–30). In the dark-tolerant cells, transcripts for all clock genes (*sasA*, *rpaA*, *kaiB*, and *kaiC*) were depleted relative to parental cells, particularly during the night (Fig. 2), indicating a potential dysregulation of the circadian rhythm, consistent with physiological measurements (Fig. 1). Circadian rhythms in cyanobacteria are known to be influenced by changes in metabolic activity (31–33), biotic interactions (34), and glucose assimilation (35,36). Thus, metabolic interactions with *Alteromonas* (7,14) could be responsible for the reduced light:dark entrainment of dark-tolerant cells (Fig. 1).

**Figure 2.**
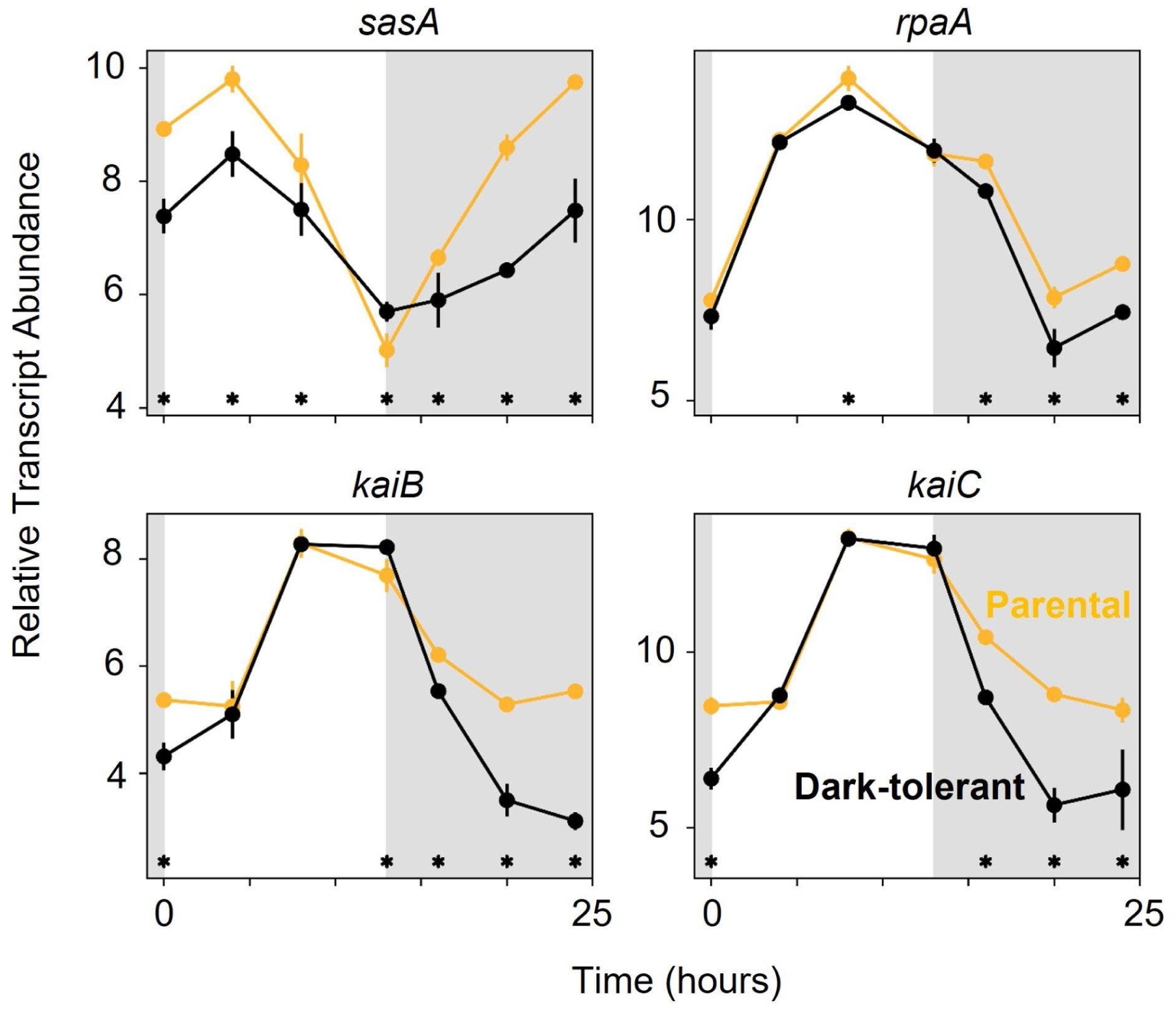
Expression patterns of the circadian clock genes of *Prochlorococcus* NATL2A, grown in co-culture with *Alteromonas*. Panels show relative transcript abundance of dark-tolerant phenotype (black line) and parental phenotype (gold line) over the 13 h day (white shading) and 11 h night (gray shading) cycle. Asterisks (*) indicate significant log2 fold differences in gene expression between the dark-tolerant and parental phenotype at the indicated time points (p <0.01).

#### Electron transport chain and ATP synthesis

We next explored how changes in the circadian rhythms of cells intersect with regulation of the electron transport chain, where the reductants and ATP that drive all metabolic processes are generated. Changes in expression were consistently seen across subunits of larger complexes in the electron transport chain (Text S2, Fig. S2). In particular, transcripts of genes involved in mediating photosynthetic electron flow (i.e., genes involved in photosystem I and II (PSI and PSII) and ferredoxin-NADP oxidoreductase (FNR) were depleted in dark-tolerant cells relative to the parental cells (Fig. 3, red panels). Conversely, transcripts of genes involved in mediating respiratory electron flow, such as NADH dehydrogenase (NDH) and cytochrome oxidase (COX), were relatively enriched (Fig. 3, blue panels), especially during the night. Transcripts of genes involved in both pathways, including cytochrome b6f (Cyt b6f) and plastocyanin (PC) showed more subtle changes (Fig. 3, black panels), but the nature of those changes were also consistent with a relative shift from autotrophy toward heterotrophy. That is, in dark-tolerant cells, transcripts for Cyt b6f and PC were relatively enriched at dusk when transcripts for respiration were most abundant, and relatively depleted in the period around dawn when transcripts for photosynthesis were most abundant. Finally, ATP synthase genes were depleted in dark-tolerant cells (Fig. 3, Table S2), suggesting a reduction in ATP synthesis. We note that one of the largest ATP-demanding processes in photosynthetic organisms is CO_2_-fixation. Taken together these data indicate a relative shift from photosynthesis toward respiration, potentially suggesting a greater use of organic carbon in dark-tolerant *Prochlorococcus*.

**Figure 3.**
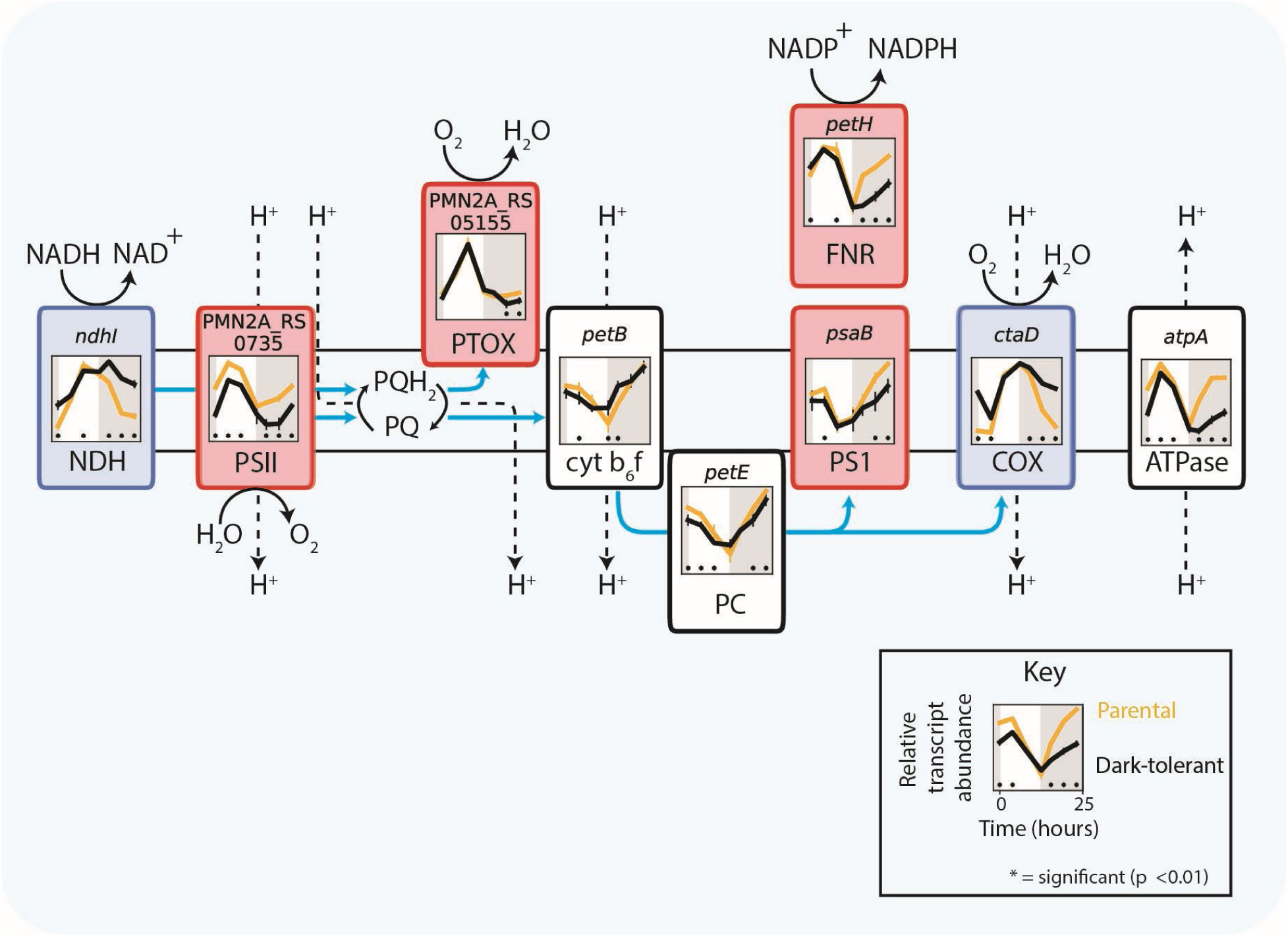
Expression patterns for genes of the photosynthetic and respiratory electron transport chain of *Prochlorococcus* NATL2A, grown in co-culture with *Alteromonas*. Panels associated with each component show relative transcript abundance of dark-tolerant (black line) and parental cultures (gold line) over the 13 h day (white shading) and 11 h night (gray shading) cycle, with representative genes for photosynthesis (red panels), respiration (blue panels), and genes involved in both pathways (black panels) (Table S2). Asterisks (*) indicate significant log2 fold differences in gene expression between the dark-tolerant and parental phenotype at the indicated time points (p <0.01). Blue arrows indicate pathways of electron flow. Abbreviations: NDH - NADH dehydrogenase, PSII - photosystem II, PQ - plastoquinone, PQH_2_ - plastoquinol, PTOX - plastoquinol terminal oxidase, cyt b6f - cytochrome b6f, PC - plastocyanin, PSI - photosystem I, FNR - ferredoxin-NADP oxidoreductase, COX - cytochrome oxidase.

#### Central carbon metabolism

Next, we examined genes involved in the central carbon metabolism of *Prochlorococcus* cells. Transcripts of genes involved in CO_2_-fixation via the Calvin cycle – a major sink of ATP and reductants – were depleted in dark-tolerant cells, whereas expression levels of genes from the pentose phosphate pathway, which feeds reductants into the respiratory electron transport chain, either changed little or increased (Fig. 4). These patterns are consistent with an increase in organic carbon use and a decrease in photosynthesis in dark-tolerant cells. Transcripts of genes that encode two enzymes in the TCA cycle, which convert fumarate and malate to oxaloacetate, were relatively depleted in dark-tolerant cells (Fig. 4), but those genes are also part of an adjacent cycle that drives a key amination step during the synthesis of ATP, which we inferred is generally reduced in dark-tolerant cells (Fig. 3). Since transcripts of other TCA cycle genes were unchanged or slightly enriched, we conclude that the respiratory function of the TCA cycle is unchanged or relatively increased in these cells. Additionally, transcripts for glycolysis genes and genes for a putative pyruvate exporter, *salY*, were depleted in dark-tolerant cells (Fig. 4). This putative decrease in pyruvate excretion, coupled to a decrease in ATP synthesis (Fig. 3) and CO_2_-fixation (Fig. 4), suggests the possibility that dark-tolerant

**Figure 4.**
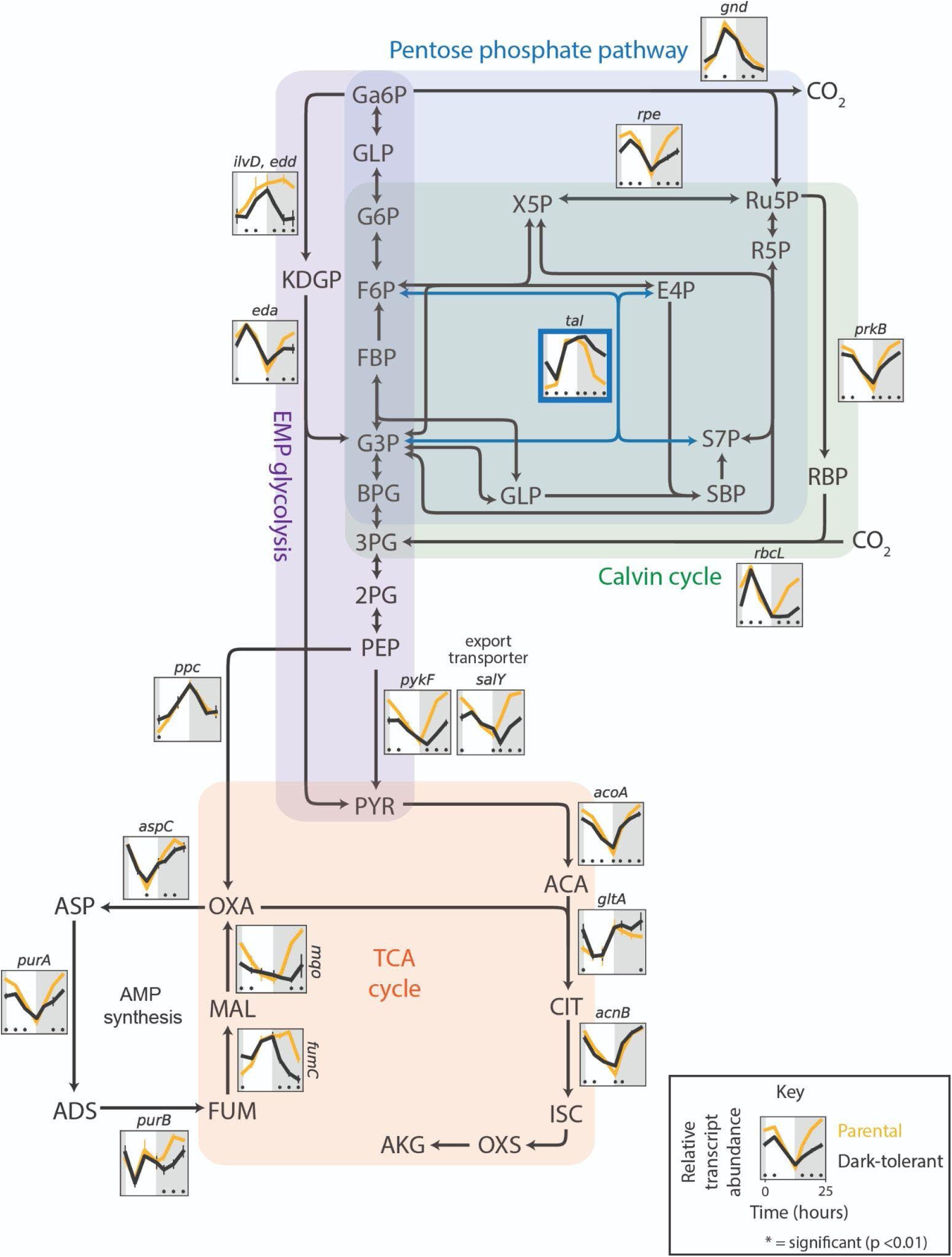
Expression patterns of genes involved in the central carbon metabolism of *Prochlorococcus* NATL2A, grown in co-culture with *Alteromonas*. The *Prochlorococcus* genes are embedded in a schematic of the Calvin cycle (green shading), glycolysis (purple shading), pentose phosphate pathway (blue shading), and TCA cycle (orange shading). Panels associated with select genes in each pathway show relative transcript abundance of dark-tolerant phenotype (black line) and parental phenotype (gold line) over the 13 h day (white shading) and 11 h night (gray shading) cycle, with representative genes shown for multi-subunit complexes (Table S3). Asterisks (*) indicate significant log2 fold differences in gene expression between the dark-tolerant and parental phenotype at the indicated time points (p <0.01). Abbreviations: Ga6P - gluconate 6-phosphate, GLP- glucono-1,5-lactono 6-phosphate, G6P – glucose 6-phosphate, F6P – fructose 6-phosphate, X5P – xylulose 5-phosphate, Ru5P – ribulose 5-phosphate, R5P – ribose 5-phosphate, E4P – erythrose 4-phosphate, S7P – sedoheptulose 7-phosphate, SBP – sedoheptulose bisphosphate, RBP – ribulose biphosphate, FBP - fructose 1,6-bisphosphate, G3P – glyceraldehyde 3- phosphate, BPG – 2,3 bisphosphoglycerate, 3PG – 3-phosphoglycerate, 2PG - 2-phosphoglycerate, PEP – phosphoenolpyruvate, PYR - pyruvate, KDGP – 2-keto-3-deoxygluconate 6-phosphate, ACA - acetoacetate, CIT - citrate, ISC - isocitrate, OXS - oxalosuccinate, AKG – alpha-ketoglutarate, OXA - oxaloacetate, MAL - malate, FUM - fumarate, ASP - aspartate, ADS- adenylosuccinate.

*Prochlorococcus* cells may exude a smaller fraction of the carbon they acquire relative to parental cells, contributing to longer dark-survival. Finally, we observed changes in nearly all regulatory genes driving these metabolic changes, hinting at a broader regulatory framework governing dark tolerance (Fig. S3, Text S3).

While expression profiles of ETC and core carbon metabolism genes are consistent with a decrease in photosynthesis and an increase in using organic carbon obtained from *Alteromonas* by dark-tolerant *Prochlorococcus*, an alternative explanation for dark-tolerance could be that cells have acquired a greater ability to store and reoxidize organic carbon to survive extended darkness. To explore this, we looked at genes related to metabolism of glycogen, considered the major carbon storage compound in *Prochlorococcus* (15). Expression for all glycogen synthesis genes is lower in dark-tolerant cells, potentially indicating a decrease in glycogen storage (Fig. 5). However, glycogen phosphorylase (GlgP), which mediates glycogen breakdown to glucose-1-phosphate, is significantly higher around sunrise (Fig. 5), which may support the transition to daytime metabolism in dark-tolerant cells, as glycogen degradation has in other cyanobacteria been found to facilitate initiation of photosynthesis by using reactions shared with the Calvin cycle (37). It has been observed that *Prochlorococcus* exudes substantial levels of polysaccharides when grown with heterotrophs (38). Hence, a decrease in expression of glycogen synthesis genes - even as expression of glycogen usage genes increase - could reflect a decrease in polysaccharide exudation, consistent with our inference from core metabolic pathways that dark-tolerant cells exude less carbon (Figs. 3 and 4 and surrounding discussion).

**Figure 5.**
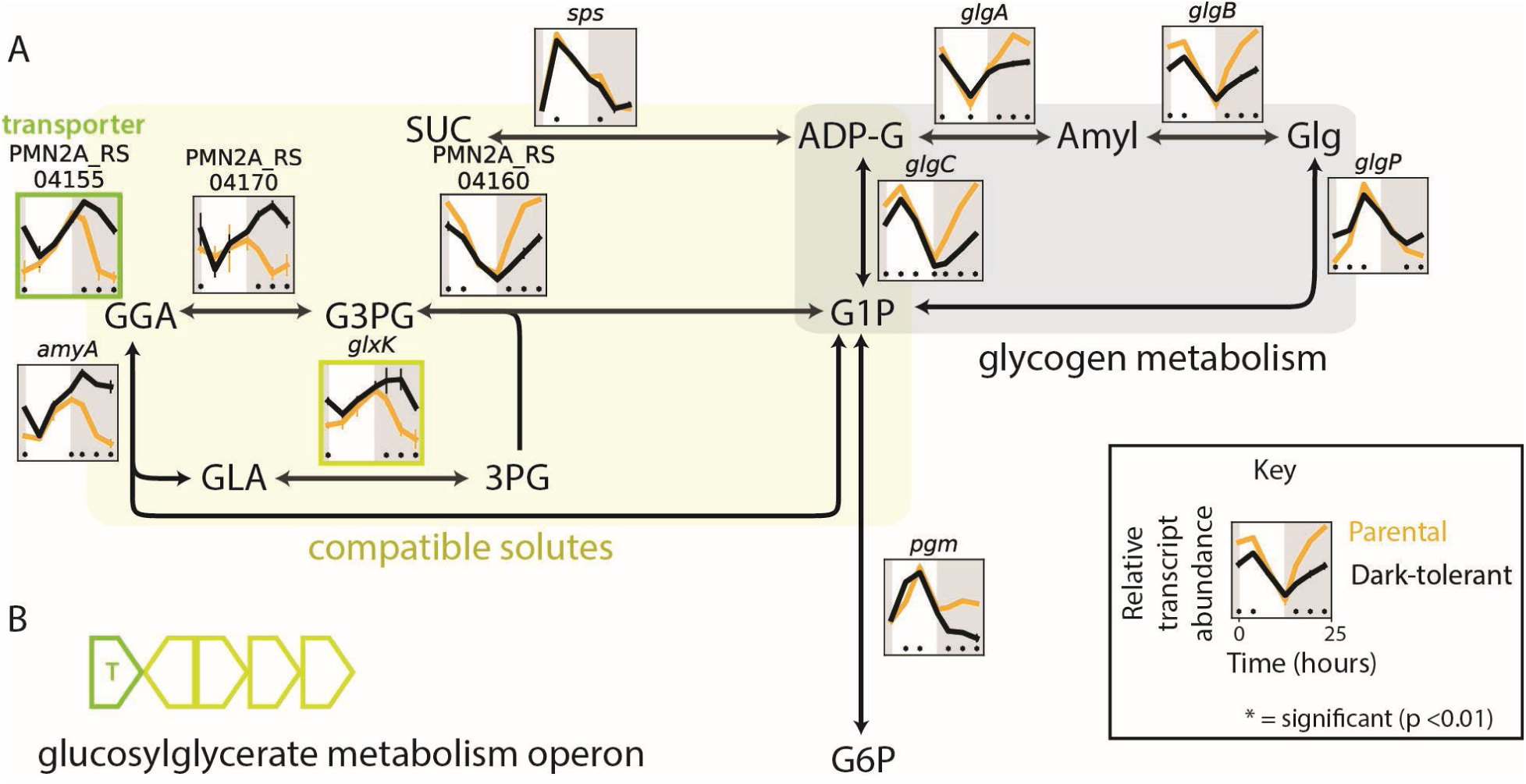
Expression patterns of glycogen metabolism and compatible solute genes in *Prochlorococcus* NATL2A, grown in co-culture with *Alteromonas*. (A) Schematic overview of metabolism of glycogen (gray shading) and compatible solutes (yellow-green shading). Panels show relative transcript abundance of dark-tolerant phenotype (black line) and parental phenotype (gold line) over a 13 h day (white shading) and 11 h night (gray shading) cycle, with representative genes shown for multi-subunit complexes (Table S2). (B) Compatible solute genes (outlined in dark yellow) are co-located in the glucosylglycerate metabolism operon and are immediately adjacent to the transporter gene (outlined in green) reported in the upper left of panel (A). Asterisks (*) indicate significant log2 fold differences in gene expression between the dark-tolerant and parental phenotype at the indicated time point (p <0.01). Abbreviations: SUC – sucrose, ADP-G – ADP-glucose, Amyl - amylose, Glg - glycogen, G1P – glucose-1-phosphate, G6P – glucose-6-phosphate, GGA- glucosylglycerate, G3PG – glucosyl 3-phosphoglycerate, 3PG – 3- phosphoglycerate, GLA – glycerate.

#### Compatible solutes

small molecules that are generally used for osmoprotection – can make up large fractions of cellular resources and could be used as an alternative energy storage mechanism in dark-tolerant *Prochlorococcus*. The major compatible solutes in *Prochlorococcus* NATL2A are sucrose and glucosylglycerate (39), the breakdown of which generates glucose-6-phosphate and/or 3-phosphoglycerate (Fig. 5), which feed directly into respiration pathways (Fig. 4). Transcripts of genes involved in sucrose metabolism were largely unchanged between dark-tolerant and parental cells, but there were clear transcriptional changes in the metabolism of glucosylglycerate (Fig. 5). In particular, transcripts of genes involved in the *de novo* synthesis of glucosyl 3-phosphoglycerate (G3PG), the immediate precursor to glucosylglycerate, were depleted in dark-tolerant cells, while those for genes involved in the recycling of glucosylglycerate to glucose-1-phosphate (G1P) and 3-phosphoglycerate (3PG) were enriched (Fig. 5). In addition, transcripts for a putative transporter gene that we identified within the glucosylglycerate metabolism operon in *Prochlorococcus* (40) are also enriched in dark-tolerant cells (Fig. 5). These patterns suggest that dark-tolerant *Prochlorococcus* is not making greater use of compatible solutes as an energy storage mechanism, but instead raise the possibility that glucosylglycerate could be an external source of organic carbon it receives from *Alteromonas*.

### Transcriptional patterns in Alteromonas

Next, we explored *Alteromonas* metabolism in the presence of both dark-adapted and parental *Prochlorococcus* cells. In contrast to the extensive transcriptional changes in *Prochlorococcus*, only 9% of *Alteromonas* genes had significant differences in expression between parental and dark-tolerant co-cultures (Fig. S4). While a large number of *Prochlorococcus* genes showed differential expression between dark-tolerant and parental phenotype at night (Fig. S1B), differential expression in *Alteromonas* was relatively more common during the daytime (Fig. S4B), suggesting *Alteromonas* may be responding to changes in metabolic byproducts released by *Prochlorococcus* during the light period.

#### Central carbon metabolism and electron transport

To explore how the emergence of dark-tolerance in *Prochlorococcus* affects *Alteromonas*, we first examined its central carbon metabolism (Fig. 6), where organic carbon obtained from *Prochlorococcus* – the only source of organic carbon for *Alteromonas* – is processed and energy is generated. While expression differences between *Alteromonas* from dark-tolerant and parental co-cultures were generally only significant at a few individual timepoints, clear trends emerged across pathways. For example, transcripts of genes in the pentose phosphate pathway, lower glycolysis and TCA cycle – the main pathways for breaking down sugars and feeding reductants into the electron transport chain – were depleted during daytime in *Alteromonas* cells from dark-tolerant relative to parental co-cultures (Fig. 6), suggesting a decrease in sugar usage and respiration during the light period. This is consistent with a concurrent decrease in the expression of Cytochrome *c* oxidase (COX) and ATP synthetase (Fig. 7), as well as the inference that dark-tolerant *Prochlorococcus* may exude less polysaccharides than parental cells (Fig. 4). An inferred decrease in the breakdown of sugars further suggests that a decrease in respiration is not linked to an increase in fermentation, which is consistent with the observation that transcripts of genes involved in producing acetate and lactate, two key fermentation products, are relatively unchanged between *Alteromonas* from dark-tolerant and parental co-cultures (Fig. S5). Collectively, these observations suggest that the increased metabolic coupling between *Prochlorococcus* and *Alteromonas* during the emergence of dark-tolerance involves a redirection of carbon flow in *Alteromonas*, likely reflecting changes in the nature of organic carbon supplied by *Prochlorococcus*.

**Figure 6.**
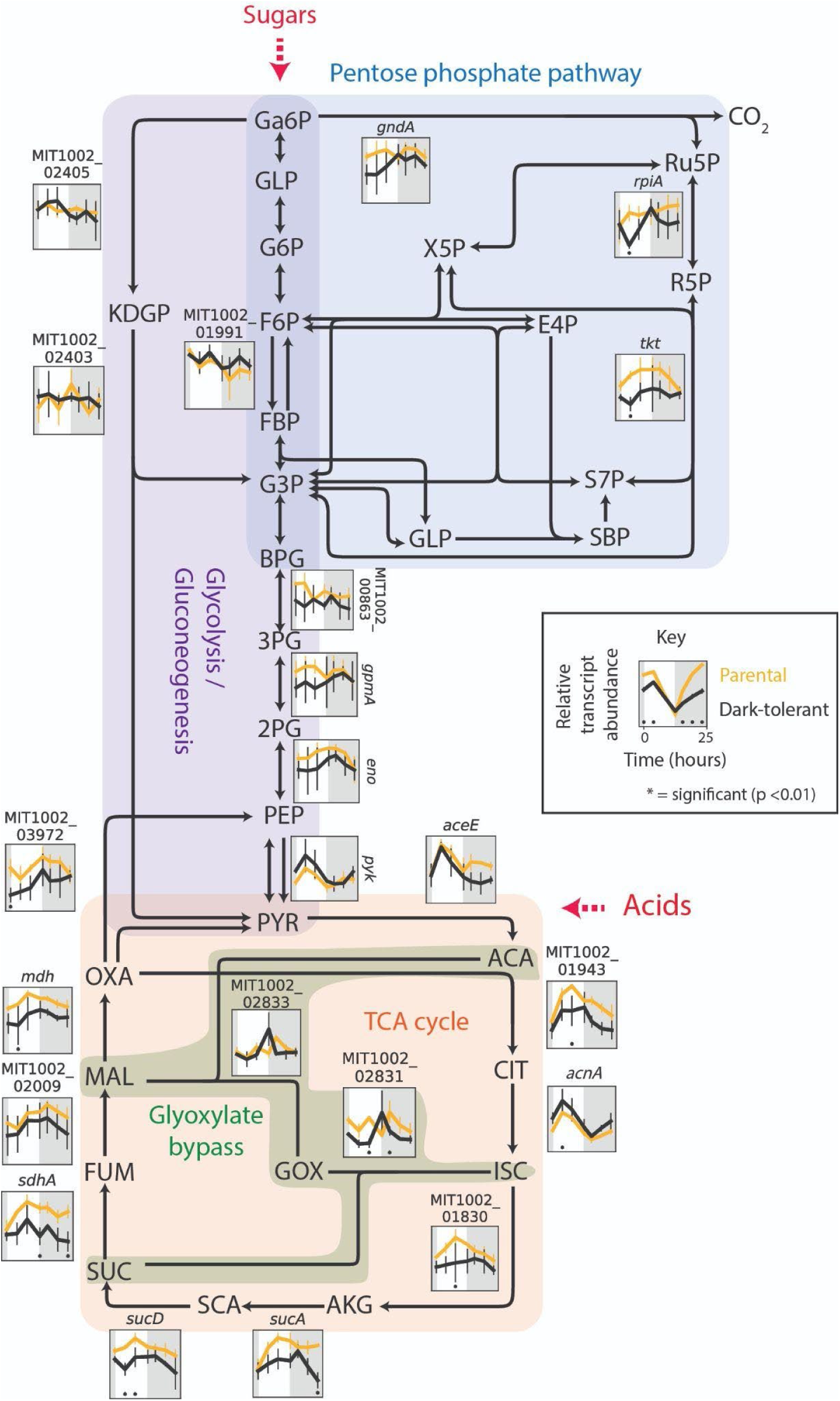
Expression patterns of genes involved in the central carbon metabolism of *Alteromonas*, grown in co-culture with *Prochlorococcus* NATL2A. Schematic of Oxidative Pentose Phosphate pathway (blue shading), ED and EMP glycolysis (purple shading), and TCA cycle (orange shading). Panels associated with select genes in each pathway show relative transcript abundance of dark-tolerant phenotype (black line) and parental phenotype (gold line) of *Alteromonas* over a 13 h day (white) and 11 h night (gray) cycle, with representative genes shown for multi-subunit complexes (Table S4). Asterisks (*) indicate significant log2 fold differences in gene expression between the dark-tolerant and parental phenotype at the indicated time point (p <0.01). Abbreviations: Ga6P - gluconate 6-phosphate, GLP- glucono-1,5- lactono 6-phosphate, G6P – glucose 6-phosphate, F6P – fructose 6-phosphate, X5P – xylulose 5- phosphate, Ru5P – ribulose 5-phosphate, R5P – ribose 5-phosphate, E4P – erythrose 4-phosphate, S7P – sedoheptulose 7-phosphate, SBP – sedoheptulose bisphosphate, RBP – ribulose biphosphate, FBP - fructose 1,6-bisphosphate, G3P – glyceraldehyde 3- phosphate, BPG – 2,3 bisphosphoglycerate, 3PG – 3- phosphoglyceric acid, 2PG - 2-phosphoglyceric acid, PEP – phosphoenolpyruvate, PYR - pyruvate, KDGP – 2-keto-3-deoxygluconate 6-phosphate, ACA - acetoacetate, CIT - citrate, ISC - isocitrate, GOX - glyoxylate, AKG – alpha-ketoglutarate, SUC - succinate, SCA - succinyl coenzyme A, OXA - oxaloacetate, MAL - malate, FUM - fumarate.

**Figure 7.**
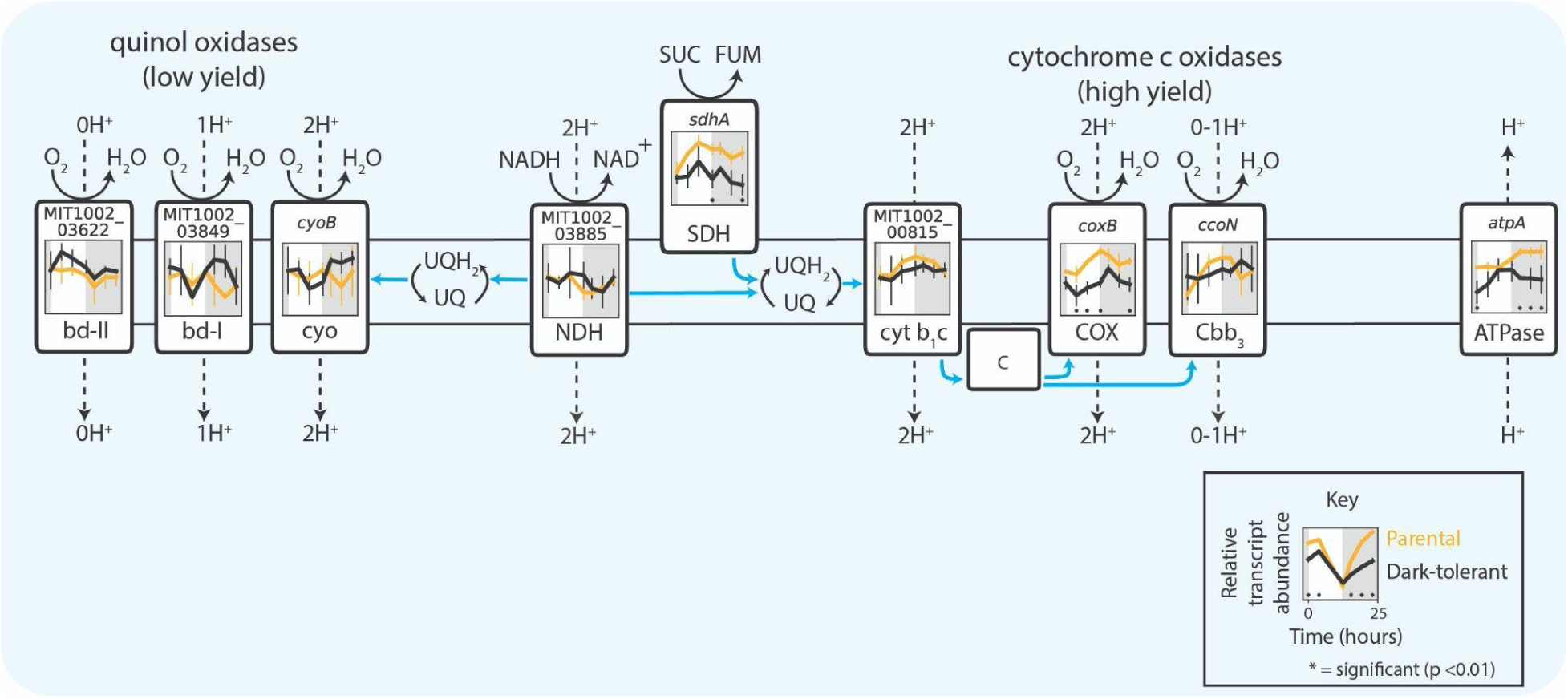
Expression patterns for genes of the electron transport chain in *Alteromonas*, grown in co-culture with *Prochlorococcus* NATL2A. The schematic includes the canonical respiration pathway, alternative terminal oxidases, and ATP synthase. Panels associated with each component show relative transcript abundance of dark-tolerant phenotype (black line) and parental phenotype (gold line) of *Alteromonas* over a 13 h day (white shading) and 11 h night (gray shading) cycle, with representative genes shown for multi-subunit complexes (Table S4). Blue arrows represent electron flow and asterisks (*) indicate significant log2 fold differences in gene expression between the dark-tolerant and parental phenotype at the indicated time point (p <0.01). Abbreviations: NDH - NADH dehydrogenase, SDH - succinate dehydrogenase, UQ - ubiquinone, QH_2_ - ubiquinol, cyt b1c - cytochrome b1c, cbb3 - cytochrome cbb3 oxidase, COX - cytochrome oxidase, bd-II and bd-I - cytochrome *bd* oxidases, cyo - cytochrome O ubiquinol oxidase.

Heterotrophic metabolism generally operates in one of two basic modes: glycolytic metabolism, based on the degradation of sugars, and gluconeogenic metabolism, based on the degradation of organic acids and/or amino acids (41,42). While we inferred a decrease in sugar degradation in *Alteromonas* (Fig. 6), we also observe a relatively unchanged expression of genes for the glyoxylate bypass, which often serves as the endpoint of organic acid degradation as it bypasses CO_2_-releasing reactions in the TCA cycle and preserves carbon backbones for gluconeogenesis and biosynthesis (43,44). Together this suggests a relative shift toward a more gluconeogenic metabolism in *Alteromonas* during the emergence of dark-tolerance in *Prochlorococcus*. Additional clues on the changing nature of organic carbon used by *Alteromonas* in dark-tolerant co-cultures come from its highly versatile electron transport chain, which can mediate multiple different pathways of electron flow that pump different numbers of protons per electron transported (Fig. 7). That is, while transcripts for cytochrome c oxidase (COX) – whose use yields the most protons pumped per electron transported – were depleted, transcripts for other lower-yield complexes were either unchanged or slightly enriched in *Alteromonas* from dark-tolerant co-cultures. This suggests that the emergence of a more gluconeogenic metabolism in *Alteromonas* is linked to a lowering of the H^+^/e^−^-pumping stoichiometry of its electron transport chain, in turn suggesting the carbon substrates it uses are more reduced in dark-tolerant co-cultures than in parental co-cultures.

### Substrate cross-feeding and the role of stress in dark-tolerant co-cultures

In an attempt to identify potential substrates from *Prochlorococcus* that fuel *Alteromonas*’ shift toward a gluconeogenic metabolism, we examined expression of *Alteromonas* genes involved in catabolism of diverse substrates (Table S5). While expression in most pathways appear relatively unchanged, we observed enrichment of transcripts for genes involved in degradation of tyrosine, phenylalanine, proline, threonine, tryptophan, benzoate, and purines in *Alteromonas* from dark-tolerant co-cultures (Fig. S6). Purines, tyrosine, hydroxybenzoate, and phenylalanine have each been identified as exudates of *Prochlorococcus* (45). Further, levels of tyrosine and phenylalanine have been shown to be elevated in cultures of *Prochlorococcus* MIT0801, which belongs to the same ecotype as NATL2A (46), compared to strains from other ecotypes (45). Moreover, transcripts of several biosynthesis genes of tyrosine, proline and purines were elevated in dark-tolerant *Prochlorococcus* cells (Fig. S7). An enrichment of transcripts involved in the biosynthesis of purines and some amino acids in *Prochlorococcus* and of transcripts for degradation of those same compounds in *Alteromonas* hints at their potential cross-feeding between the two species.

Kujawinski et al (2023) identified a putative link between cellular stress and amino acid exudation in *Prochlorococcus*, suggesting that stress could play a role in reinforcing putative cross-feeding of amino acids in dark-tolerant co-cultures. This is consistent with observations that under stress many other bacteria, including other cyanobacteria, activate the stringent response pathway, a regulatory mechanism designed to slow down growth and conserve cellular resources (47–49), in some cases specifically to drive an increase in amino acid biosynthesis (50). Similarly, we observe significant changes in multiple general stress and stringent response genes in dark-tolerant cells compared to parental *Prochlorococcus* cells (Fig. S8, Text S4), including nighttime enrichment of transcripts for the hibernation promoting factor *hpf* (Fig. S8D) in dark-tolerant cells, which binds to ribosomes, reducing ribosomal activity and protein synthesis to conserve energy (48,51). Collective observations suggest that the stress from a lack of energy due to extended darkness may trigger the stringent response in *Prochlorococcus*, which then drives an increase in amino acid synthesis, thereby changing the nature of cross-feeding with *Alteromonas*.

## Conclusions

This work suggests that the mechanisms underlying dark-tolerance in *Prochlorococcus* co-cultured with *Alteromonas* involve a dynamic interplay of their metabolisms (Fig. 8). The observed changes in gene expression suggest a model in which metabolic adjustments of both partners shape the flow of carbon and energy between the two taxa (Fig. 8). Dark-tolerant *Prochlorococcus* cells showed a loosened coupling between cell division and the light:dark cycle (Fig. 1), a reduction in transcripts involved in photosynthesis and an enrichment of transcripts involved in respiration (Fig. 3), which point at a relative shift from photosynthesis toward respiration throughout the diel light:dark cycle (Fig. 8, yellow arrows). Further, dark-tolerant *Prochlorococcus* showed a lower growth rate and its gene expression patterns indicate a reduction in CO_2_-fixation and ATP synthesis (Fig. 3 and 4) and a putative decrease in excretion of pyruvate and possibly polysaccharides in comparison to parental cells, hinting at a more economical use of energy and organic carbon (Fig. 8, red arrows). These physiological (Fig. 1) and metabolic (Fig. 3-5) adaptations underscore the multifaceted nature of dark-tolerance, which is driven by broad-scale adjustments in multiple regulatory mechanisms (Fig. 2 and S3).

**Figure 8.**
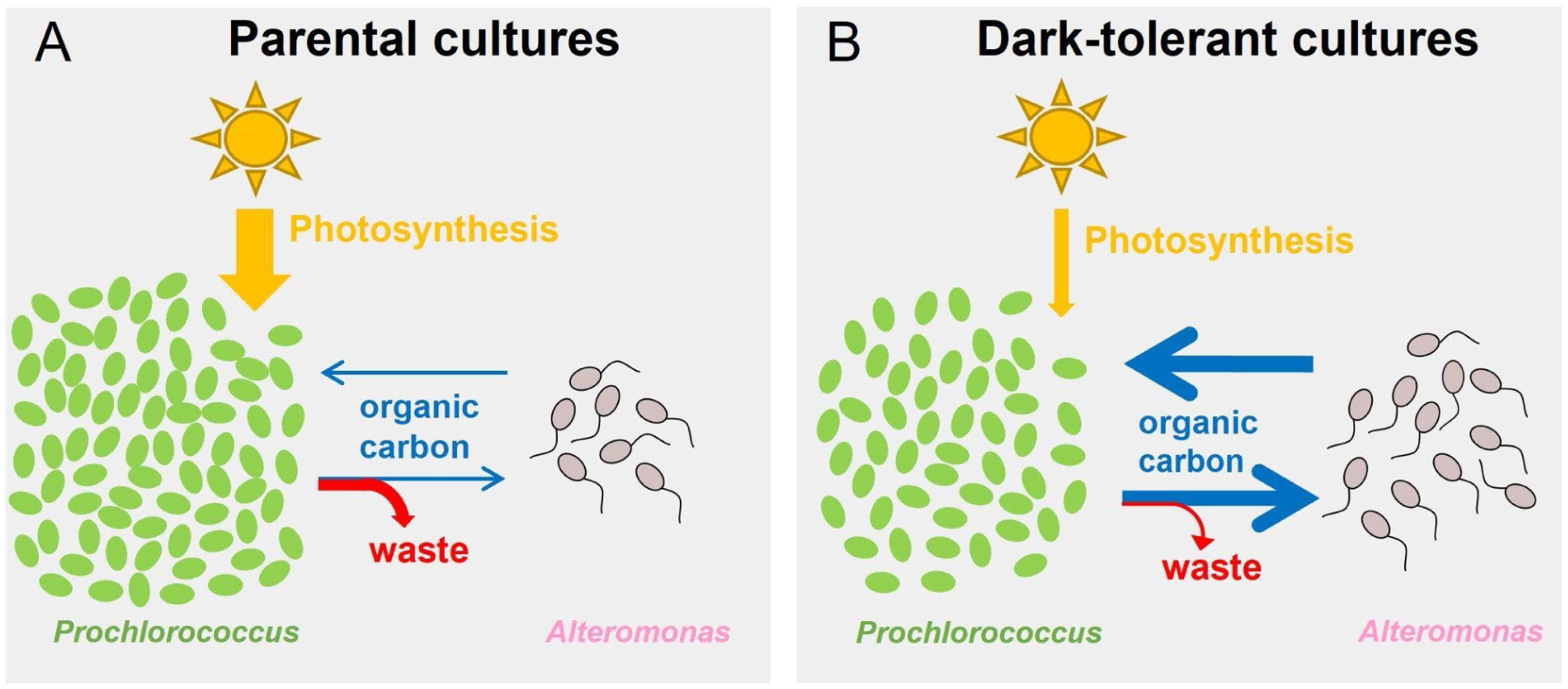
Summary of inferred changes in the metabolic coupling between *Prochlorococcus* and *Alteromonas* associated with the emergence of dark-tolerance. (A) *Prochlorococcus* parental cells rely heavily on photosynthesis (yellow arrow), have less carbon exchange (blue arrows) with *Alteromonas*, and waste more metabolic products (red arrow) that are not assimilated by *Alteromonas*. (B) Dark-tolerant *Prochlorococcus* show a relative shift from photosynthetic to respiratory processes, a more economical use of energy and organic carbon (i.e., they produce less “waste”), and a greater mutual exchange of organic carbon with *Alteromonas* (blue arrows). The stronger metabolic coupling between the two species results in a higher *Alteromonas*:*Prochlorococcus* cell ratio.

As shown by our previous studies (7), extended dark survival of *Prochlorococcus* depends on the presence of a heterotroph, such as *Alteromonas*. Conversely, the medium supporting the co-cultures of *Prochlorococcus* and *Alteromonas* lacks supplemented organic carbon sources, and therefore the heterotroph *Alteromonas* is fully dependent on *Prochlorococcus* for its carbon supply. The increased *Alteromonas*:*Prochlorococcus* ratio in dark-tolerant cultures (Fig. 1), the tighter coupling of their population dynamics, and the implied relative shift toward heterotrophic metabolism in *Prochlorococcus*, suggests that this metabolic coupling is strengthened compared to parental co-cultures due to an enhanced carbon exchange between the two partners (Fig. 8, blue arrows). Our results hint at the organic carbon compounds that might be involved. The gene expression patterns in *Alteromonas* cells from dark-tolerant co-cultures indicate a reduction in use of respiratory pathways, a decrease in the H^+^/e^-^ pumping stoichiometry, and a shift away from using sugars towards more reduced organic acids, likely reflecting changes in the nature of organic carbon supplied by *Prochlorococcus* (Fig. 6 and 7). In line with this, the data showed an enrichment of transcripts involved in the biosynthesis of amino acids and purines in dark-tolerant *Prochlorococcus* (Fig. S7) and an enrichment of transcripts for degradation of those same compounds in *Alteromonas* (Fig. S6). We speculate that, in return, the release of some partially processed organic carbon by *Alteromonas* could be a way to maintain its redox balance, and at the same time, provide *Prochlorococcus* with the organic carbon needed to survive extended darkness (Fig. 8, blue arrows). Indeed, we observed a reduction in transcripts for biosynthesis, combined with an increase of transcripts for the degradation and transport, of the compatible solute glucoslyglycerate in *Prochlorococcus* (Fig. 5), suggesting that dark-tolerant *Prochlorococcus* cells might be acquiring this organic carbon source from *Alteromonas* (Fig. 8, blue arrows).

The results of this study lead us to suspect that in the wild, *Prochlorococcus* is able to survive prolonged periods in the dark waters of the deep ocean through adaptations that promote mutual exchange of carbon and energy with surrounding microbial cells with complementary metabolisms.

## Acknowledgements

The authors thank both past and present members of the Chisholm Lab for their support and comments, especially Sean Kearney and Christine Ziegler for their valuable input. This study was funded by the Simons Foundation (Life Sciences Project Award ID 337262, S.W.C.; Life Sciences Project Award ID 509034SCFY20, R.B., S.W.C; Life Sciences SCOPE Award ID 329108, S.W.C.; SCOPE Award ID 721246; S.W.C; Life Sciences Award ID 917971, S.J.B.) and the National Science Foundation (OCE-2019589, R.B.; OCE-2049004 and OCE-2304066, S.J.B.)

## Data Availability

All sequencing data is available through NCBI Gene Expression Omnibus, GEO series accession number GSE264347. Supplemental Table data is available through figshare, doi: 10.6084/m9.figshare.25727190. Data from physiological studies and bacterial strains are available by request from the corresponding authors.

## References

1. Flombaum P, Gallegos JL, Gordillo RA, Rincón J, Zabala LL, Jiao N, et al. Present and future global distributions of the marine Cyanobacteria *Prochlorococcus* and *Synechococcus*. PNAS. 2013;110(24):9824–9.

2. Johnson ZI, Zinser ER, Coe A, McNulty NP, Woodward EMS, Chisholm SW. Niche partitioning among *Prochlorococcus* ecotypes along ocean-scale environmental gradients. Science. 2006;311(5768):1737–40.

3. Tseng CH, Chiang PW, Lai HC, Shiah FK, Hsu TC, Chen YL, et al. Prokaryotic assemblages and metagenomes in pelagic zones of the South China Sea. BMC Genomics. 2015;16(1):219.

4. Sebastián M, Giner CR, Balagué V, Gómez-Letona M, Massana R, Logares R, et al. The active free-living bathypelagic microbiome is largely dominated by rare surface taxa. ISME Communications. 2024;4(1):ycae015.

5. Mende DR, Boeuf D, DeLong EF. Persistent Core Populations Shape the Microbiome Throughout the Water Column in the North Pacific Subtropical Gyre. Front Microbiol. 2019;10.

6. Jiao N, Luo T, Zhang R, Yan W, Lin Y, Johnson ZI, et al. Presence of *Prochlorococcus* in the aphotic waters of the western Pacific Ocean. Biogeosciences. 2014;11(8):2391–400.

7. Coe A, Ghizzoni J, LeGault K, Biller S, Roggensack SE, Chisholm SW. Survival of *Prochlorococcus* in extended darkness. Limnology and Oceanography. 2016;61(4):1375–88.

8. Zubkov MV, Tarran GA, Fuchs BM. Depth related amino acid uptake by *Prochlorococcus* cyanobacteria in the Southern Atlantic tropical gyre. FEMS Microbiol Ecol. 2004;50(3):153– 61.

9. Vila-Costa M, Simó R, Harada H, Gasol JM, Slezak D, Kiene RP. Dimethylsulfoniopropionate Uptake by Marine Phytoplankton. Science. 2006;314(5799):652– 4.

10. Michelou VK, Cottrell MT, Kirchman DL. Light-Stimulated Bacterial Production and Amino Acid Assimilation by Cyanobacteria and Other Microbes in the North Atlantic Ocean. Appl Environ Microbiol. 2007;73(17):5539–46.

11. Mary I, Tarran GA, Warwick PE, Terry MJ, Scanlan DJ, Burkill PH, et al. Light enhanced amino acid uptake by dominant bacterioplankton groups in surface waters of the Atlantic Ocean. FEMS Microbiol Ecol. 2008;63(1):36–45.

12. Muñoz-Marín M del C, Luque I, Zubkov MV, Hill PG, Diez J, García-Fernández JM. *Prochlorococcus* can use the Pro1404 transporter to take up glucose at nanomolar concentrations in the Atlantic Ocean. Proc Natl Acad Sci USA. 2013;110(21):8597–602.

13. Wu Z, Aharonovich D, Roth-Rosenberg D, Weissberg O, Luzzatto-Knaan T, Vogts A, et al. Single-cell measurements and modelling reveal substantial organic carbon acquisition by *Prochlorococcus*. Nat Microbiol. 2022;7(12):2068–77.

14. Coe A, Biller SJ, Thomas E, Boulias K, Bliem C, Arellano A, et al. Coping with darkness: The adaptive response of marine picocyanobacteria to repeated light energy deprivation. Limnology and Oceanography. 2021;66(9):3300–12.

15. Zinser ER, Lindell D, Johnson ZI, Futschik ME, Steglich C, Coleman ML, et al. Choreography of the transcriptome, photophysiology, and cell cycle of a minimal photoautotroph, *Prochlorococcus*. PLoS ONE. 2009;4(4):e5135.

16. Steglich C, Futschik M, Rector T, Steen R, Chisholm SW. Genome-Wide Analysis of Light Sensing in *Prochlorococcus*. Journal of Bacteriology. 2006;188(22):7796–806.

17. Biller SJ, Coe A, Chisholm SW. Torn apart and reunited: impact of a heterotroph on the transcriptome of *Prochlorococcus*. ISME J. 2016;10(12):2831–43.

18. Bushnell B. BBTools Software Package. 2014. https://sourceforge.net/projects/bbmap/

19. Anders S, Pyl PT, Huber W. HTSeq—a Python framework to work with high-throughput sequencing data. Bioinformatics. 2015;31(2):166–9.

20. Love MI, Huber W, Anders S. Moderated estimation of fold change and dispersion for RNA-seq data with DESeq2. Genome Biology. 2014;15(12):550.

21. Wickham H. ggplot2. WIREs Computational Statistics. 2011;3(2):180–5.

22. Thaben PF, Westermark PO. Detecting rhythms in time series with RAIN. J Biol Rhythms. 2014;29(6):391–400.

23. Waldbauer JR, Rodrigue S, Coleman ML, Chisholm SW. Transcriptome and proteome dynamics of a light-dark synchronized bacterial cell cycle. PLoS ONE. 2012;7(8):e43432.

24. Holtzendorff J, Partensky F, Mella D, Lennon JF, Hess WR, Garczarek L. Genome streamlining results in loss of robustness of the circadian clock in the marine cyanobacterium *Prochlorococcus marinus* PCC 9511. J Biol Rhythms. 2008;23(3):187–99.

25. Axmann IM, Dühring U, Seeliger L, Arnold A, Vanselow JT, Kramer A, et al. Biochemical Evidence for a Timing Mechanism in *Prochlorococcus*. Journal of Bacteriology. 2009;191(17):5342–7.

26. Diamond S, Rubin BE, Shultzaberger RK, Chen Y, Barber CD, Golden SS. Redox crisis underlies conditional light–dark lethality in cyanobacterial mutants that lack the circadian regulator, RpaA. Proceedings of the National Academy of Sciences. 2017;114(4):E580–9.

27. Iwasaki H, Williams SB, Kitayama Y, Ishiura M, Golden SS, Kondo T. A kaiC-interacting sensory histidine kinase, SasA, necessary to sustain robust circadian oscillation in cyanobacteria. Cell. 2000;101(2):223–33.

28. Smith RM, Williams SB. Circadian rhythms in gene transcription imparted by chromosome compaction in the cyanobacterium *Synechococcus elongatus*. Proceedings of the National Academy of Sciences. 2006;103(22):8564–9.

29. Takai N, Nakajima M, Oyama T, Kito R, Sugita C, Sugita M, et al. A KaiC-associating SasA–RpaA two-component regulatory system as a major circadian timing mediator in cyanobacteria. Proc Natl Acad Sci U S A. 2006;103(32):12109–14.

30. Dong G, Yang Q, Wang Q, Kim YI, Wood TL, Osteryoung KW, et al. Elevated ATPase Activity of KaiC Applies a Circadian Checkpoint on Cell Division in *Synechococcus elongatus*. Cell. 2010;140(4):529–39.

31. Golden SS. Timekeeping in bacteria: the cyanobacterial circadian clock. Curr Opin Microbiol. 2003;6(6):535–40.

32. Pattanayak GK, Phong C, Rust MJ. Rhythms in Energy Storage Control the Ability of the Cyanobacterial Circadian Clock to Reset. Current Biology. 2014;24(16):1934–8.

33. Pattanayak GK, Lambert G, Bernat K, Rust MJ. Controlling the Cyanobacterial Clock by Synthetically Rewiring Metabolism. Cell Reports. 2015;13(11):2362–7.

34. Biller SJ, Coe A, Roggensack SE, Chisholm SW. Heterotroph Interactions Alter *Prochlorococcus* Transcriptome Dynamics during Extended Periods of Darkness. mSystems. 2018;3(3):e00040–18.

35. Muñoz-Marín MDC, Duhamel S, Björkman KM, Magasin JD, Díez J, Karl DM, et al. Differential Timing for Glucose Assimilation in Prochlorococcus and Coexistent Microbial Populations in the North Pacific Subtropical Gyre. Microbiol Spectr. 2022;e0246622.

36. Moreno-Cabezuelo JÁ, Gómez-Baena G, Díez J, García-Fernández JM. Integrated Proteomic and Metabolomic Analyses Show Differential Effects of Glucose Availability in Marine *Synechococcus* and *Prochlorococcus*. Microbiology Spectrum. 2023;0(0):e03275–22.

37. Shinde S, Zhang X, Singapuri SP, Kalra I, Liu X, Morgan-Kiss RM, et al. Glycogen Metabolism Supports Photosynthesis Start through the Oxidative Pentose Phosphate Pathway in Cyanobacteria. Plant Physiol. 2020;182(1):507–17.

38. Cruz BN, Neuer S. Heterotrophic Bacteria Enhance the Aggregation of the Marine Picocyanobacteria *Prochlorococcus* and *Synechococcus*. Front Microbiol. 2019;10.

39. Klähn S, Steglich C, Hess WR, Hagemann M. Glucosylglycerate: a secondary compatible solute common to marine cyanobacteria from nitrogen-poor environments. Environ Microbiol. 2010;12(1):83–94.

40. Franceus J, Pinel D, Desmet T. Glucosylglycerate Phosphorylase, an Enzyme with Novel Specificity Involved in Compatible Solute Metabolism. Appl Environ Microbiol. 2017;83(19):e01434–17.

41. Schink SJ, Christodoulou D, Mukherjee A, Athaide E, Brunner V, Fuhrer T, et al. Glycolysis/gluconeogenesis specialization in microbes is driven by biochemical constraints of flux sensing. Mol Syst Biol. 2022;18(1):e10704.

42. Gralka M, Pollak S, Cordero OX. Genome content predicts the carbon catabolic preferences of heterotrophic bacteria. Nat Microbiol. 2023;8(10):1799–808.

43. Kornberg HL. The role and control of the glyoxylate cycle in Escherichia coli. Biochem J. 1966;99(1):1–11.

44. Cronan, JE, Laporte D. Tricarboxylic Acid Cycle and Glyoxylate Bypass. EcoSal Plus. 2005;1(2).

45. Kujawinski EB, Braakman R, Longnecker K, Becker JW, Chisholm SW, Dooley K, et al. Metabolite diversity among representatives of divergent *Prochlorococcus* ecotypes. mSystems. 2023;8(5):e01261–22.

46. Biller SJ, Berube PM, Berta-Thompson JW, Kelly L, Roggensack SE, Awad L, et al. Genomes of diverse isolates of the marine cyanobacterium *Prochlorococcus*. Sci Data. 2014;1:140034.

47. Wells DH, Gaynor EC. Helicobacter pylori Initiates the Stringent Response upon Nutrient and pH Downshift. Journal of Bacteriology. 2006;188(10):3726–9.

48. Hood RD, Higgins SA, Flamholz A, Nichols RJ, Savage DF. The stringent response regulates adaptation to darkness in the cyanobacterium *Synechococcus elongatus*. PNAS. 2016;113(33):E4867–76.

49. Calderón-Flores A, Du Pont G, Huerta-Saquero A, Merchant-Larios H, Servín-González L, Durán S. The Stringent Response Is Required for Amino Acid and Nitrate Utilization, Nod Factor Regulation, Nodulation, and Nitrogen Fixation in *Rhizobium etli*. J Bacteriol. 2005;187(15):5075–83.

50. Paul BJ, Berkmen MB, Gourse RL. DksA potentiates direct activation of amino acid promoters by ppGpp. Proc Natl Acad Sci U S A. 2005;102(22):7823–8.

51. Prossliner T, Gerdes K, Sørensen MA, Winther KS. Hibernation factors directly block ribonucleases from entering the ribosome in response to starvation. Nucleic Acids Research. 2021 Feb 26;49(4):2226–39.

