## Supplemental Text 1-4, Tables 1-6, Figures 1-8 for "Emergence of metabolic coupling to the heterotroph *Alteromonas* promotes dark survival in *Prochlorococcus*"

### Methods

### RNA concentration

First pass RNA yields were low, so a technical replicate RNA sample was extracted and then combined and concentrated with the first technical replicate using a re-precipitation method. To the combined RNA, a 10% volume of 2.73M pH 5.2 sodium acetate and 4°C 100% ethanol was added along with 1 µl of glycogen and mixed. The samples were placed into -80°C freezer overnight to allow for precipitation. The following day, the samples were centrifuged for 35 min at 12,000 *x g* and 4°C to collect the precipitate. The supernatant was carefully removed and precipitated RNA was washed twice with 4°C 75% ethanol, spinning for 10 min at 12,000 *x g* and 4°C. After the second wash, the remaining ethanol was carefully removed and discarded by pipet and the RNA was left to dry for up to 10 min at room temperature. The RNA was dissolved in 25µl 10mM pH 7.0 Tris and stored at -80 °C.

**Cell Cycle Periodicity**

Clusters of co-expressed genes were identified using the soft R clustering package Mfuzz (1). The optimal number of clusters was chosen based on evaluation of overlap, inertia plot, and cluster dissimilarity plots. Clear separation of gene expression patterns is reflected in the absence of overlap among clusters, which, in turn, indicates a favorable number of clusters. In addition, the selection of the cluster number was guided by inertia plot and cluster dissimilarity. Minimal drops in both signify coherent partitions of gene clusters while minimizing redundancy between clusters.

While the periodicity in this experiment is higher than the 69% reported in a previous study using the same co-culture (2), it is more similar to data found with another *Prochlorococcus* strain, MED4, which showed 94% of genes exhibited periodicity (3). Both of these studies sampled more frequently over the diel and used the CYCLE algorithm, but due to the decreased number of timepoints in our study, we found the algorithms used in previous experiments were not as effective in determining periodicity as using RAIN with this dataset.

**Text 1**

*Transcriptional patterns and periodicity in Prochlorococcus*

To obtain an understanding of the metabolic changes associated with the dark-tolerant phenotype, we compared the transcriptomes of parental and dark-tolerant cells using RNA-Seq (Table S3). Gene expression patterns among *Prochlorococcus* biological replicates were tightly correlated in both parental and dark-tolerant cells across the diel cycle (Fig. S1 A), which highlights the higher degree of similarity in gene expression patterns among biological replicates. The vast majority of *Prochlorococcus* genes (97%) were significantly differentially expressed between the dark-tolerant and parental strains for at least one time point during the diel cycle (Fig. S1 B, Table S2). This is consistent with widespread changes to cell physiology (Fig. 1). More genes showed significantly differential expression in the dark-tolerant compared to the parental cells during nighttime than during daytime (median absolute log2 fold change of 0.75±0.7 at night vs 0.47±0.36 in the day; Fig. S1 C), suggesting a greater impact on cellular metabolism and regulation during periods of darkness.

Because the coupling of the cell cycle with the light:dark cycle was relaxed in the dark-tolerant cells compared to the parental cells (Fig. 1), we expected to see a similar decoupling effect in the diel gene expression patterns. To our surprise, we found that 94% of the *Prochlorococcus* transcriptome was expressed with significant periodicity in both the parental and dark-tolerant cells (Table S2). However, while dark-tolerant gene expression patterns remained periodic overall, only 31% of genes were expressed with the same periodicity pattern as seen in the parental cells (Table S2), likely attributed to changes in gene expression observed during the night (Fig. S1 C). These data suggest that differential diel regulation of gene expression timing and/or amplitude likely contribute to the profound impact that dark adaptation has on overall gene expression.

**Text 2**

*Genes related to photosynthesis and ATP synthesis in Prochlorococcus*

As we observed reduced entrainment to the light:dark cycle in dark-tolerant *Prochlorococcus*, we wondered how cellular metabolism and energy production were affected by these changes. To examine this, we explored the response to genes involved in the electron transport chain. We observed a decrease in transcripts for photosynthesis and ATP synthase coupled with an increase in transcripts involved in respiration in dark-tolerant cells relative to parental cells (Fig. 3). Additionally, genes encoding prochlorophyte chlorophyll-binding proteins (Pcb), which bind divinyl chlorophylls *a* and *b* and are part of the light harvesting complex in *Prochlorococcus* (4,5), showed similar patterns as the genes involved in photosynthetic electron transport. In fact, transcripts for all 7 *pcb* genes identified in NATL2A were depleted in dark-tolerant relative to parental cells (Fig. S2), which is consistent with the shift away from photosynthesis (Figs. 1 and 3). Similarly, expression of cytochrome M (*cytM*), a gene thought to play a role in regulating photosynthetic capacity under mixotrophic conditions in *Synechocystis* (6,7), was depleted in dark-tolerant relative to parental culture (Fig. S2). These data hint at a metabolic shift from photosynthesis to respiration.

Transcripts for AtpΘ, a small protein encoded by the gene *atpT* in cyanobacteria responsible for preventing the reverse reaction (hydrolysis) of ATP synthase during low-energy conditions (i.e. night or extended darkness) (8), were significantly enriched in dark-tolerant cells relative to parental cells during the night (Fig. S2). We hypothesize that dark-tolerant cells, characterized by reduced ATP synthesis (Fig. 3), increase *atpT* expression during the night aiming to safeguard against ATP loss in anticipation of prolonged darkness.

**Text 3**

*Regulatory genes in Prochlorococcus*

We wanted to explore potential gene expression mechanisms behind the decrease in photosynthesis and increase organic carbon use in dark-tolerant *Prochlorococcus* (Fig. 3 and 4). To examine this, we investigated regulatory genes including transcription factors, sigma factors, and small RNAs (sRNAs). We observed significant differences in nearly all regulatory genes in dark-tolerant cells compared to parental cells (Fig. S3), suggesting profound alterations in the regulatory landscape of these cells in response to dark adaptation. Notably, we found significant changes in 18 of the 19 putative transcription factors (9,10) in dark-tolerant cells relative to the parental cells (Fig. S3). Specifically, transcripts for ribonucleotide reductase, *nrdR*, and ferric uptake regulator, *fur*, transcription repressors that regulate ribonucleotide reductase and metabolism, respectively, were enriched in dark-tolerant cells relative to parental cells during the night (Fig. S3). Overexpression of *nrdR* causes reduced bacterial growth, fitness, and adherence characteristics in *E.coli* (11), which could explain the reduced growth rates observed in dark-tolerant cells (Fig. 1). Next, we found that transcripts for 4 of the 5 five sigma factors found in *Prochlorococcus* NATL2A (9) were enriched in dark-tolerant cells relative to the parental cells during the night (Fig. S3). All four of these genes are considered group I sigma factors, which transcribe housekeeping and basic cellular function genes and are thought to be essential for cell viability.

Finally, we examined small RNAs (sRNAs), for which most have unknown functions, but are important for transcriptional and post-transcriptional regulation of gene expression in *Prochlorococcus* (12,13). These sRNAs are particularly relevant under conditions of environmental stresses, such as light adaptation, nutrient limitation, and phage infection (14,15), and may be essential for adaptive physiological and metabolic adaptations (16,17). Although our libraries were not specifically prepared for enrichment of sRNAs, we identified reads mapping to Yfr genes, which are relatively long sRNA sequences. Transcripts for 15 of the 18 Yfr genes were significantly differentially abundant at one or more time points, with Yfr1, Yfr2, Yfr13, Yfr23, Yfr103, Yfr107, and Yfr108 showing more pronounced changes in dark-tolerant cultures relative to parental cultures (Fig. S3, Table S2). Lambrecht et al (2019) examined the function of Yfr1 and Yfr2 in *Prochlorococcus* MED4 and found that these motifs are complementary to each other, suggesting an interdependence in their regulatory network and discovered these genes have hundreds of mRNA targets belonging to functional classes such as translation, transcription, and DNA replication (i.e. repair for Yfr1 and photosynthesis, respiration, and regulatory functions for Yfr2). In our study, Yfr1 exhibits ~ 2x reduction in expression during the night in dark-tolerant cells compared to parental cells (Fig. S3) and conversely, in one of the four copies of Yfr2, the opposite trend is observed (Fig. S3), aligning with findings in Lambrecht et al, 2019. Finally, we found that transcripts for two copies of sRNA Yfr103 were more abundantly expressed in dark-tolerant cells than the parental cells during the night (Fig. S3). Yfr103 has been implicated in the response to fluctuating light, temperature, and darkness in both laboratory and natural *Prochlorococcus* populations (18,19), and is the most abundant sRNA in the ocean (20,21). When combined with our data this suggests that Yfr103 has an important regulatory function in *Prochlorococcus*, possibly in dark-adaptation.

Data presented here suggests that multiple overlapping regulatory mechanisms influence dark adaptation. While it is not entirely clear why the expression of most of these proteins changes only during the night in dark-tolerant cells, it is possible the cells could be optimizing metabolic processes in order to allocate resources efficiently to ensure survival in the absence of light or "priming" metabolic processes in anticipation of returning to the light after prolonged darkness.

**Text 4**

*Genes related to general stress response in Prochlorococcus*

Physiology and metabolic data suggest dark-tolerant *Prochlorococcus* cells had a slower growth rate (Fig. 1), underwent metabolic shifts involving less photosynthesis (Fig. 3), and enhanced amino acid synthesis (Fig. S7). As Kujawinski et al (2023) found a possible relationship between stress and amino acid exudation, we wondered whether these cells were stressed. To examine this, we explored changes to general stress response and stringent response pathway genes. Shifts in general stress response genes such as *clpB*, *dnaJ*, *groES* were evident in the transcriptomes of dark-tolerant cells (Fig. S8). Transcripts for *groES* and *dnaJ*, both of which are involved in protein folding and preventing misfolding (22), were depleted at almost every time point in dark-tolerant cells, with the exception of three time points where transcripts for *groES* were enriched (Fig. S8). As protein folding is a high-energy function requiring ATP and dark-tolerant *Prochlorococcus* cells are possibly making less ATP (Fig. 3), these data indicate a general disruption to or a shift away from allocating resources towards protein folding. Transcripts for *clpB*, which is involved in disaggregation of proteins (23), were depleted near sunset and enriched during the night in dark-tolerant cells compared to parental cells (Fig. S8). This hints at possible increased protein aggregation either during the regular night period or in anticipation of prolonged darkness. Combined, these data suggest a shift in how dark-tolerant *Prochlorococcus* cells maintain protein homeostasis.

Another regulatory system bacteria can utilize in response to stressful conditions (i.e nutrient or energy limitation) is the stringent response pathway (24). In an earlier study examining *Prochlorococcus'* response to extended darkness, we found that only 16 out of the 42 stringent response genes (homologous to those found in *Synechococcus elongatus* PCC7942) were differentially expressed (2). As results from the previous study examined an immediate or "shock" response to extended darkness, it made us wonder how the "long-term" adaptation in this study would change expression patterns in this pathway. Notably, 41 out of 42 stringent response genes were significantly differentially expressed at at least one time point in dark-tolerant relative to parental cells (Supplemental Table 3). Additionally, 21 out of the 42 stringent response genes were differentially expressed at all night time points in dark-tolerant cells relative to parental. Specifically, we observed nighttime enrichment of transcripts for *spoT* (Fig. S8E), which synthesizes guanosine tetraphosphate or pentaphosphate ((p)ppGpp) (25), are also enriched in dark-tolerant *Prochlorococcus*. When subjected to darkness, freshwater *Synechococcus elongatus* accumulates (p)ppGpp, resulting in reduced cell growth, translation, and potentially decreased pigment production (26). These data and previous findings suggest the stringent response could be important in *Prochlorococcus'* dark survival by slowing down growth (Fig. 1), reducing photosynthetic activity (Fig. 1 and 3), and maintaining translational machinery, which, in turn, could lead to the observed increase in expression of amino acid biosynthesis genes (Fig. S7).

### References


1. Kumar L, E Futschik M. Mfuzz: a software package for soft clustering of microarray data. Bioinformation. 2007;2(1):5–7.

2. Biller SJ, Coe A, Roggensack SE, Chisholm SW. Heterotroph Interactions Alter *Prochlorococcus* Transcriptome Dynamics during Extended Periods of Darkness. mSystems. 2018;3(3):e00040-18.

3. Zinser ER, Lindell D, Johnson ZI, Futschik ME, Steglich C, Coleman ML, et al. Choreography of the transcriptome, photophysiology, and cell cycle of a minimal photoautotroph, *Prochlorococcus*. PLoS ONE. 2009;4(4):e5135.

4. Bibby TS, Mary I, Nield J, Partensky F, Barber J. Low-light-adapted *Prochlorococcus* species possess specific antennae for each photosystem. Nature. 2003;424(6952):1051–4.

5. Garczarek L, Dufresne A, Rousvoal S, West NJ, Mazard S, Marie D, et al. High vertical and low horizontal diversity of *Prochlorococcus* ecotypes in the Mediterranean Sea in summer. FEMS Microbiol Ecol. 2007;60(2):189–206.

6. Solymosi D, Nikkanen L, Muth-Pawlak D, Fitzpatrick D, Vasudevan R, Howe CJ, et al. Cytochrome cM Decreases Photosynthesis under Photomixotrophy in *Synechocystis* sp. PCC 68031 [CC-BY]. Plant Physiology. 2020;183(2):700–16.

7. Hiraide Y, Oshima K, Fujisawa T, Uesaka K, Hirose Y, Tsujimoto R, et al. Loss of cytochrome cM stimulates cyanobacterial heterotrophic growth in the dark. Plant Cell Physiol. 2015;56(2):334–45.

8. Song K, Hagemann M, Georg J, Maaß S, Becher D, Hess WR. Expression of the Cyanobacterial FoF1 ATP Synthase Regulator AtpΘ Depends on Small DNA-Binding Proteins and Differential mRNA Stability. Microbiology Spectrum. 2022 Apr 21;10(3):e02562-21.

9. Kiełbasa SM, Herzel H, Axmann IM. Regulatory elements of marine cyanobacteria. Genome Inform. 2007;18:1–11.

10. Lambrecht SJ, Steglich C, Hess WR. A minimum set of regulators to thrive in the ocean. FEMS Microbiology Reviews. 2020;44(2):232–52.

11. Naveen V, Hsiao CD. NrdR Transcription Regulation: Global Proteome Analysis and Its Role in *Escherichia coli* Viability and Virulence. PLoS One. 2016;11(6):e0157165.

12. Steglich C, Lindell D, Futschik M, Rector T, Steen R, Chisholm SW. Short RNA half-lives in the slow-growing marine cyanobacterium *Prochlorococcus*. Genome Biol. 2010;11(5):R54.

13. Raghavan R, Groisman E, Ochman H. Genome-wide detection of novel regulatory RNAs in *E. coli*. https://genome-cshlp-org.libproxy.mit.edu/content/21/9/1487.short

14. Steglich C, Futschik ME, Lindell D, Voss B, Chisholm SW, Hess WR. The challenge of regulation in a minimal photoautotroph: non-coding RNAs in *Prochlorococcus*. PLoS Genet. 2008;4(8):e1000173.

15. Lambrecht SJ, Kanesaki Y, Fuss J, Huettel B, Reinhardt R, Steglich C. Interplay and Targetome of the Two Conserved Cyanobacterial sRNAs Yfr1 and Yfr2 in *Prochlorococcus* MED4. Scientific Reports. 2019;9(1):14331.

16. De Porcellinis A, Frigaard NU, Sakuragi Y. Determination of the Glycogen Content in Cyanobacteria. J Vis Exp. 2017;(125):56068.

17. Breaker RR, Harris KA, Lyon SE, Wencker FDR, Fernando CM. Evidence that OLE RNA is a component of a major stress-responsive ribonucleoprotein particle in extremophilic bacteria. Molecular Microbiology. 2023;120(3):324–40.

18. Voigt K, Sharma CM, Mitschke J, Joke Lambrecht S, Voß B, Hess WR, et al. Comparative transcriptomics of two environmentally relevant cyanobacteria reveals unexpected transcriptome diversity. The ISME Journal. 2014;8(10):2056–68.

19. Miller D, Pfreundt U, Hou S, Lott S, Hess W, Berman-Frank I. Winter mixing impacts gene expression in marine microbial populations in the Gulf of Aqaba. Aquat Microb Ecol. 2017;80(3):223–42.

20. Pfreundt U, Spungin D, Bonnet S, Berman-Frank I, Hess WR. Global analysis of gene expression dynamics within the marine microbial community during the VAHINE mesocosm experiment in the southwest Pacific. Biogeosciences. 2016;13(14):4135–49.

21. Hou S, Pfreundt U, Miller D, Berman-Frank I, Hess WR. mdRNA-Seq analysis of marine microbial communities from the northern Red Sea. Sci Rep. 2016;6:35470.

22. Susin MF, Baldini RL, Gueiros-Filho F, Gomes SL. GroES/GroEL and DnaK/DnaJ Have Distinct Roles in Stress Responses and during Cell Cycle Progression in *Caulobacter crescentus*. J Bacteriol. 2006;188(23):8044–53.

23. Alam A, Bröms JE, Kumar R, Sjöstedt A. The Role of ClpB in Bacterial Stress Responses and Virulence. Frontiers in Molecular Biosciences. 2021;8.

24. Cashel M, Gallant J. Two compounds implicated in the function of the RC gene of *Escherichia coli*. Nature. 1969;221(5183):838–41.

25. Ronneau S, Hallez R. Make and break the alarmone: regulation of (p)ppGpp synthetase/hydrolase enzymes in bacteria. FEMS Microbiol Rev. 2019;43(4):389–400.

26. Hood RD, Higgins SA, Flamholz A, Nichols RJ, Savage DF. The stringent response regulates adaptation to darkness in the cyanobacterium *Synechococcus elongatus*. PNAS. 2016;113(33):E4867–76.

**Supplemental Table 1**. RNA-seq statistics for both *Prochlorococcus* and *Alteromonas*. Total library size, reads mapped to either *Prochlorococcus* NATL2A or *Alteromonas macleodii* MIT1002 genomes, and transcripts that are aligned to each ORF (sense orientation).

**Supplemental Table 2**. Significantly differentially abundant *Prochlorococcus* transcripts. Values represent log2 fold change for every time point. Statistically significantly differential transcript abundances are designated by "***" ($p < 0.01$).

**Supplemental Table 3**. Gene periodicity patterns in *Prochlorococcus* cells. Based on the periodicity pattern, genes in parental and dark-tolerant cells were clustered into 15 clusters. If a gene in both parental and dark-tolerant cells was found in the same cluster, the pattern was marked "true", and if not, as "false".

**Supplemental Table 4**. Significantly differentially abundant *Alteromonas* transcripts. Values represent log2 fold change for every time point. Statistically significantly differential transcript abundances are designated by "***" ($p < 0.01$).

**Supplemental Table 5**. Differential abundance of *Alteromonas* transcripts involved in diverse substrate catabolism. Values represent log2 fold change for every time point. Statistically significantly differential transcript abundances are designated by "***" ($p < 0.01$).

**Supplemental Table 6.** *Prochlorococcus* transcripts found to be significantly differentially expressed at every time point. Values represent log2 fold change for every time point. Statistically significantly differential transcript abundances are designated by "***" ($p < 0.01$).

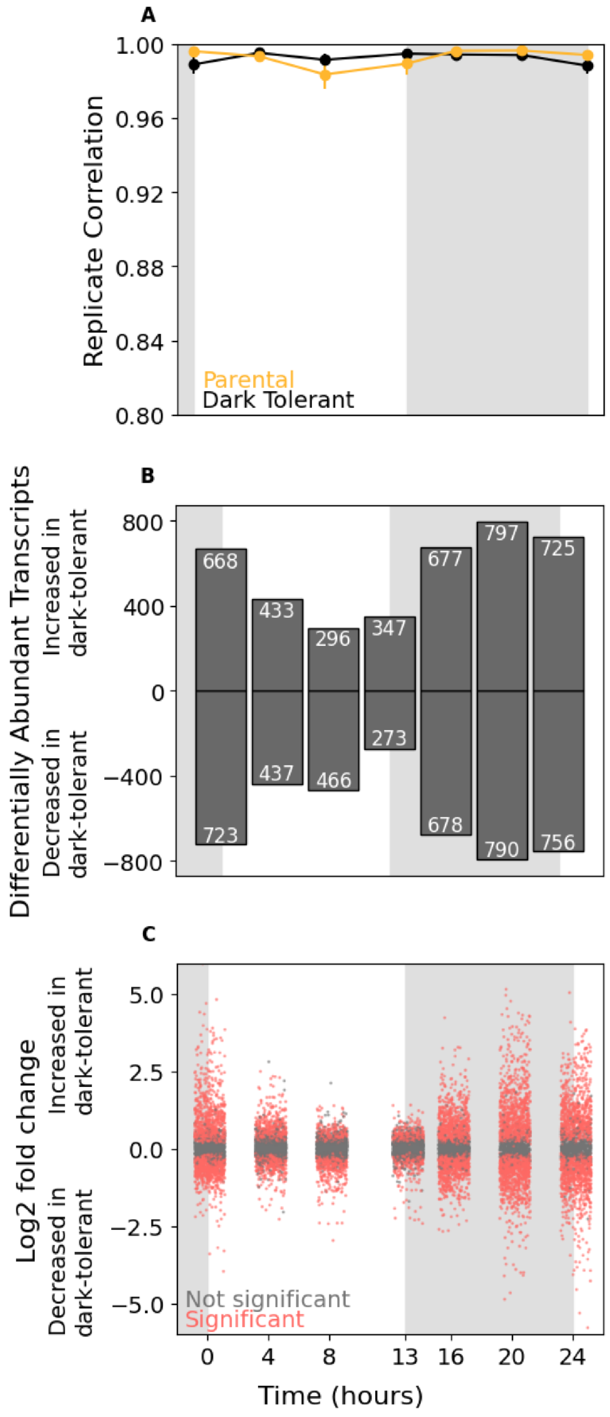


**Supplemental Figure 1**. Transcriptome changes after repeated light energy deprivation. *Prochlorococcus* NATL2A parental (gold) and dark-tolerant (black) replicate correlation (A), differentially abundant transcripts (white text) represented above (enriched) or below (depleted) the center black line (B), and log2 fold change of significant (red) and not significant transcripts (gray, C) of dark-tolerant relative to parental cells over a 24 h period while growing in 13:11 light:dark conditions. The gray shaded area indicates a standard 11 h night period.

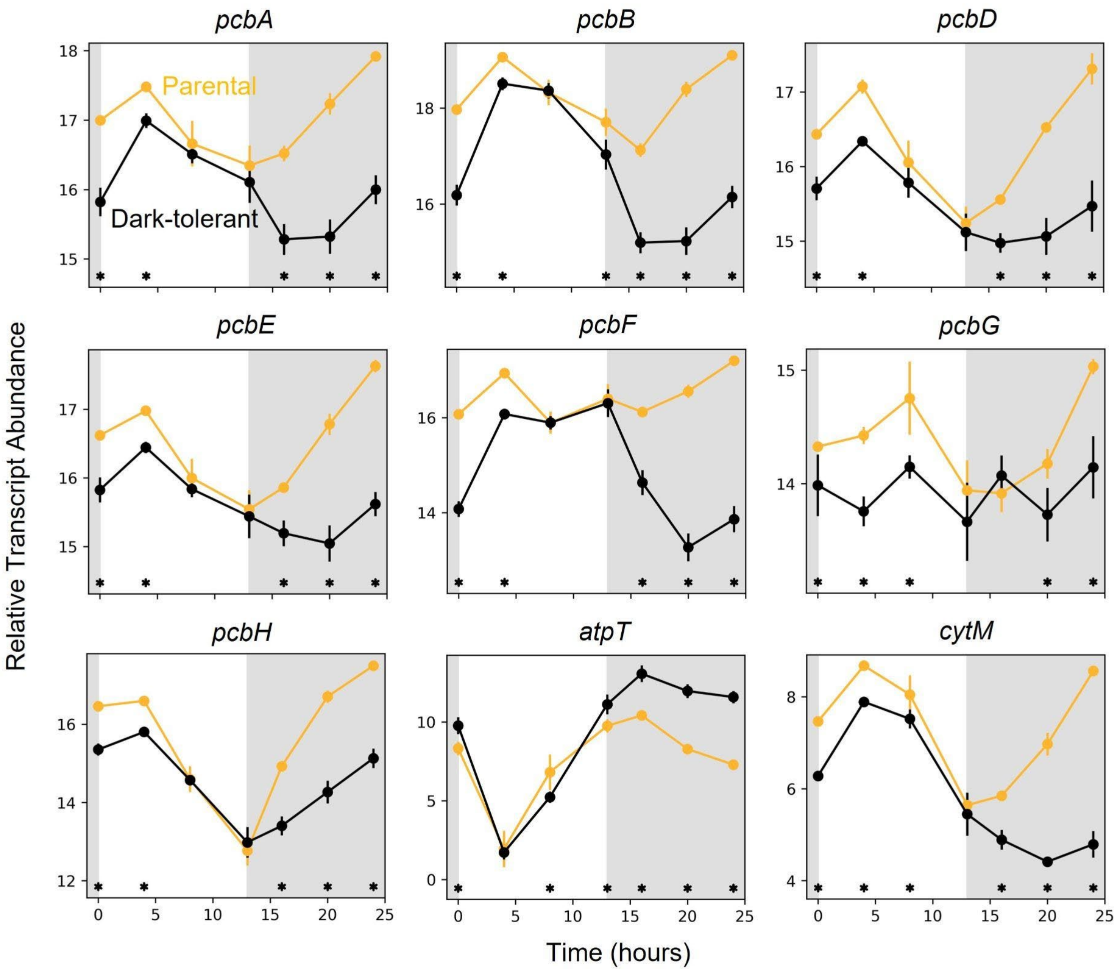


**Supplemental Figure 2.** Relative transcript abundance for genes related to photosynthesis and ATP synthesis in *Prochlorococcus* NATL2A parental (gold) and dark-tolerant cells (black) over a 13 h day (white) and 11 h night (gray) cycle. Asterisks (*) indicate significant log2 fold differences in gene expression between the dark-tolerant and parental phenotype at the indicated time point ($p < 0.01$).

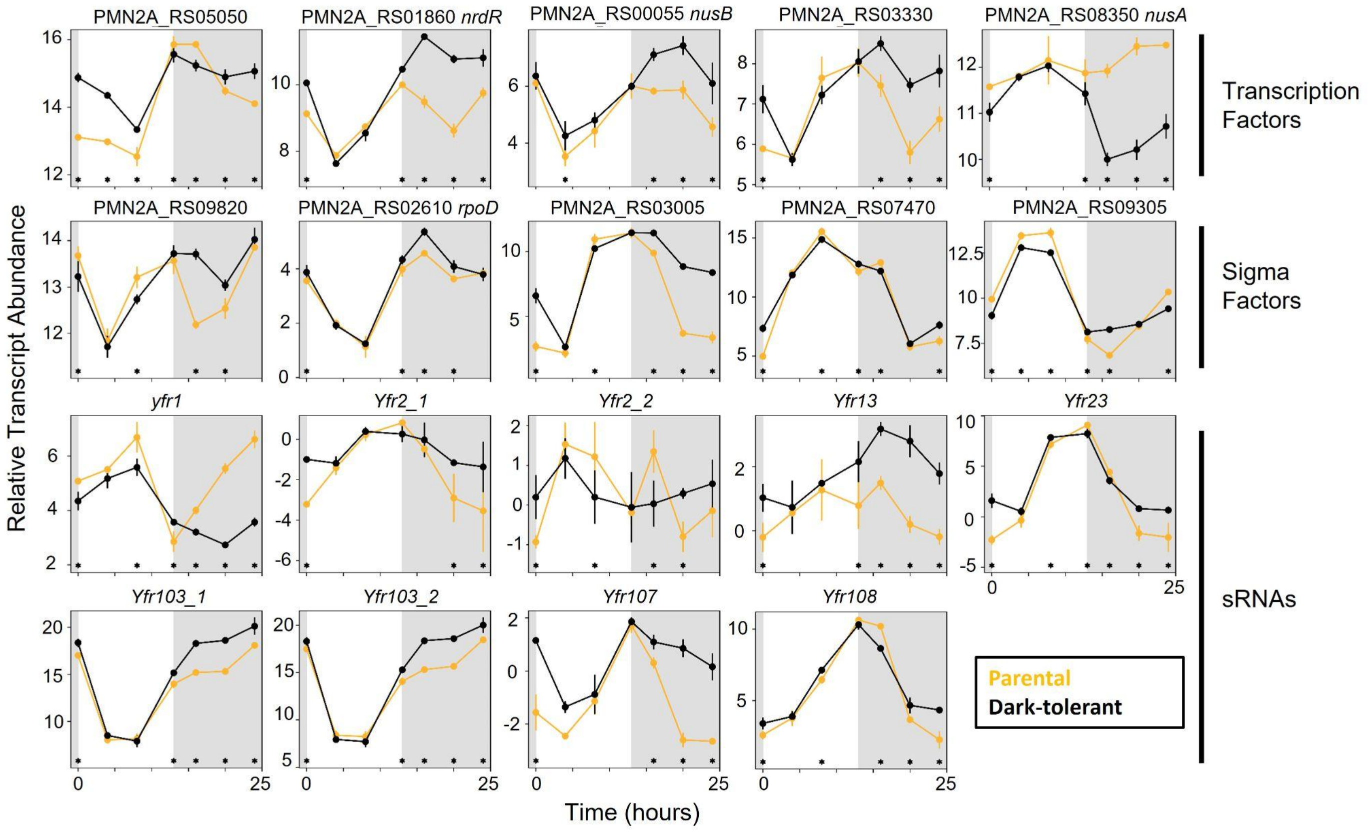


**Supplemental Figure 3.** *Prochlorococcus* regulatory protein expression patterns. Relative transcript abundance of regulatory genes, including transcription factors (A-E), sigma factors (F-J), and small RNAs (sRNAs) (K-S) in parental (gold) and dark-tolerant phenotype (black) *Prochlorococcus* NATL2A shown over a 13 h day (white) and 11 h night (gray) cycle. Asterisks (*) indicate significant $\log_2$ fold differences in gene expression between the dark-tolerant and parental phenotype at the indicated time point ($p < 0.01$). Only a select number of regulatory genes are shown, for all refer to Table S2.

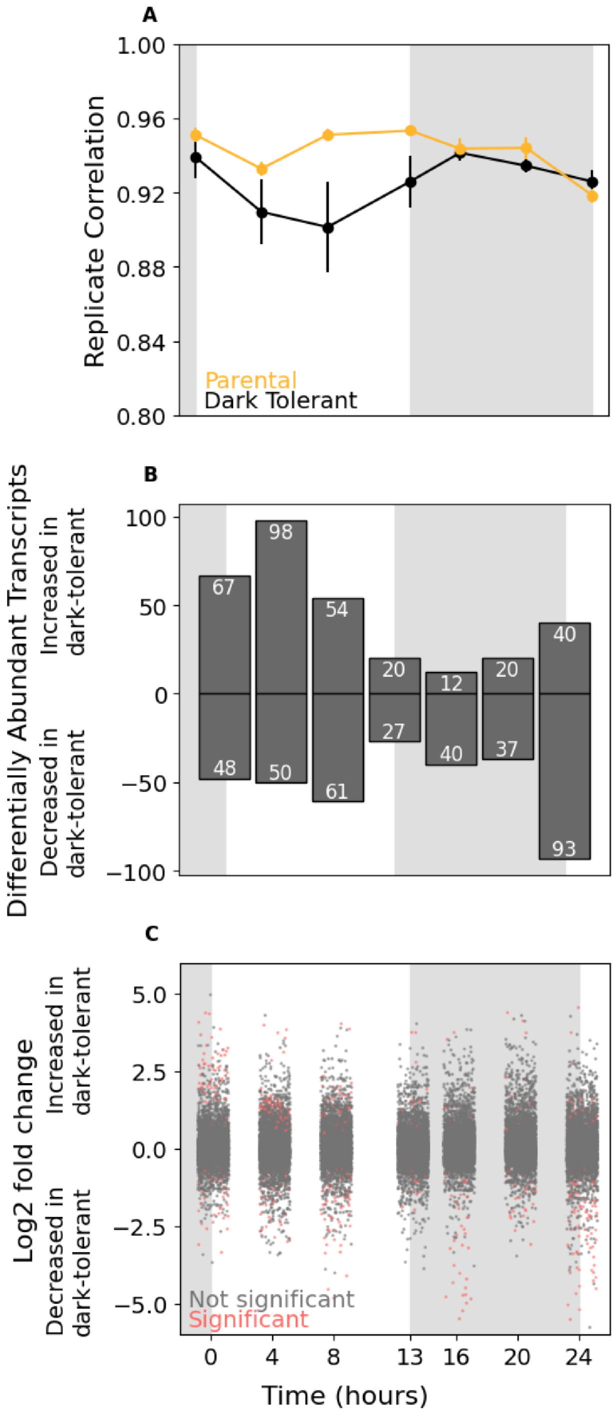


**Supplemental Figure 4**. Transcriptome changes after repeated light energy deprivation. *Alteromonas* MIT1002 replicate correlation of dark-tolerant (black) relative to parental cells (gold, no extended darkness) (A), Log2 fold change of significant (red) and not significant transcripts (gray, C), and differentially abundant transcripts (white text) represented above (enriched) or below (depleted) the center black line (B) over a 24 h period while growing in 13:11 h light:dark conditions. The gray shaded area indicates a standard 11 h night period.

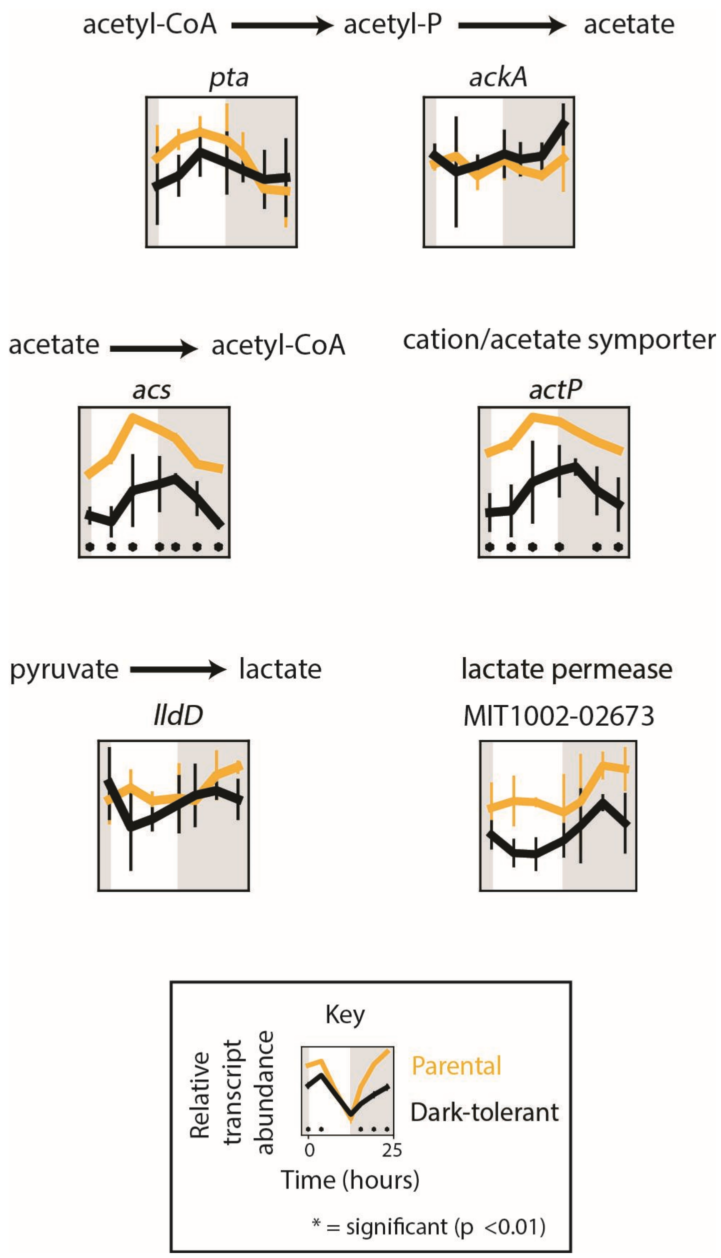


**Supplemental Figure 5**. Pathways associated with fermentation in *Alteromonas*. Acetate synthesis, acetyl-CoA synthesis, lactate synthesis, and lactate transport. Relative transcript abundance of parental (gold) and dark-tolerant phenotype (black) *Alteromonas* in co-culture with *Prochlorococcus* NATL2A shown over a 13 h day (white) and 11 h night (gray) cycle. Asterisks (*) indicate significant log2 fold differences in gene expression between the dark-tolerant and parental phenotype at the indicated time point (p <0.01).

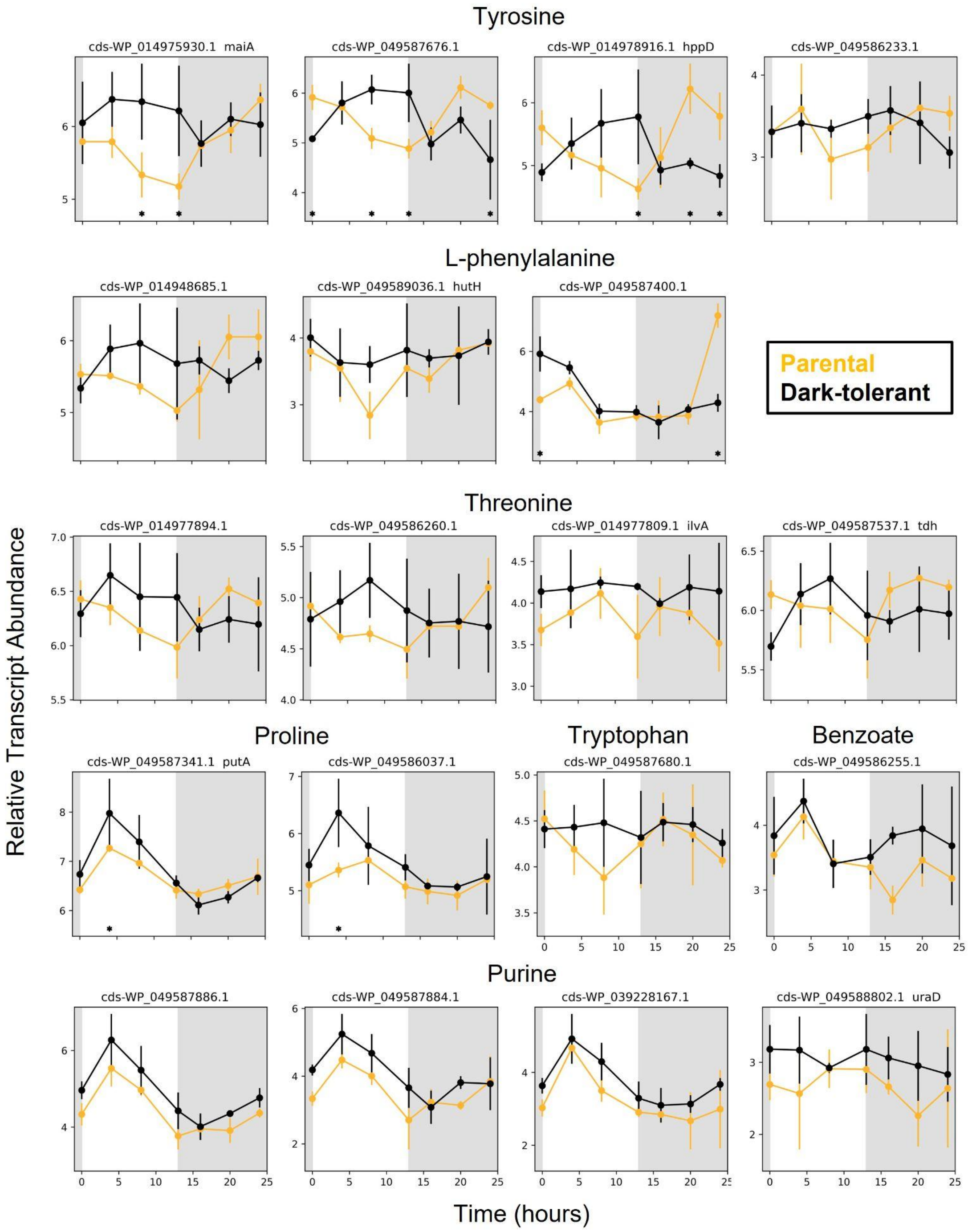


**Supplemental Figure 6**. *Alteromonas* genes associated with tyrosine, L-phenylalanine, threonine, proline, tryptophan, benzoate, and purine degradation pathways. Relative transcript abundance of parental (gold) and dark-tolerant phenotype (black) *Alteromonas* in co-culture with *Prochlorococcus* NATL2A shown over a 13 h day (white) and 11 h night (gray) cycle with representative genes shown for multi-subunit complexes (Table S6). Asterisks (*) indicate significant $\log_2$ fold differences in gene expression between the dark-tolerant and parental phenotype at the indicated time point ($p < 0.01$).

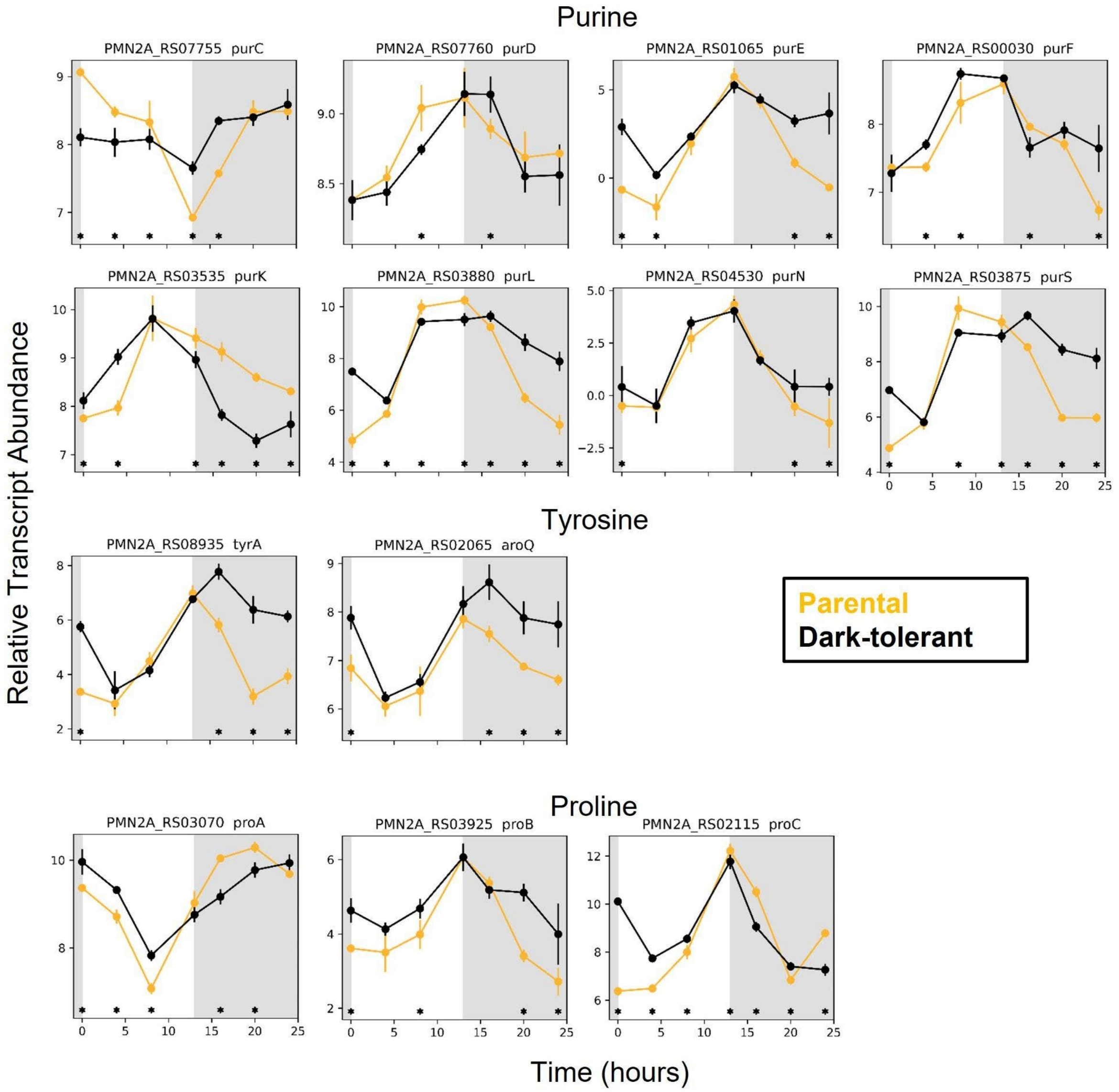


**Supplemental Figure 7.** *Prochlorococcus* purine, tyrosine, and proline biosynthesis genes. Relative transcript abundance of parental (gold) and dark-tolerant phenotype (black) *Prochlorococcus* NATL2A shown over a 13 h day (white) and 11 h night (gray) cycle with representative genes shown for multi-subunit complexes (Table S2). Asterisks (*) indicate significant log2 fold differences in gene expression between the dark-tolerant and parental phenotype at the indicated time point (p <0.01).

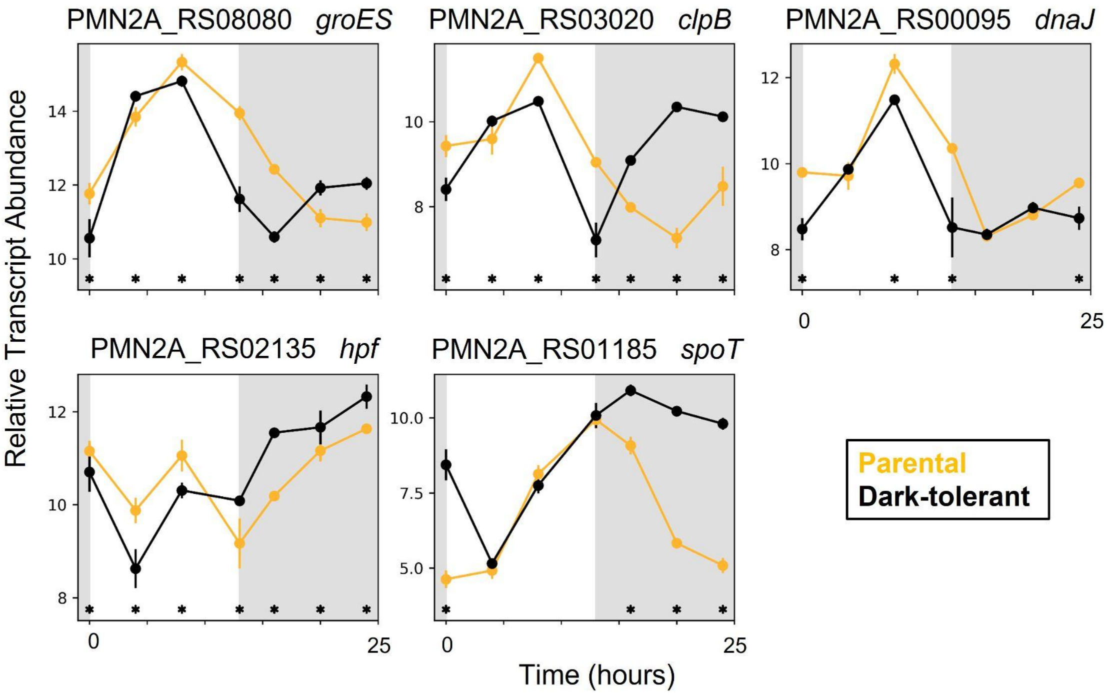


**Supplemental Figure 8**. Stress and stringent response genes in *Prochlorococcus* NATL2A. Relative transcript abundance of parental (gold) and dark-tolerant phenotype (black) *Prochlorococcus* NATL2A shown over a 13 h day (white) and 11 h night (gray) cycle. Additional stringent response genes are found in Table S2. Asterisks (*) indicate significant $\log_2$ fold differences in gene expression between the dark-tolerant and parental phenotype at the indicated time point ($p < 0.01$).
